# CytoMAP: a spatial analysis toolbox reveals features of myeloid cell organization in lymphoid tissues

**DOI:** 10.1101/769877

**Authors:** Caleb R Stoltzfus, Jakub Filipek, Benjamin H Gern, Brandy E Olin, Joseph M Leal, Miranda R Lyons-Cohen, Jessica Huang, Clarissa L Paz-Stoltzfus, Courtney R Plumlee, Thomas Pöschinger, Kevin B Urdahl, Mario Perro, Michael Y Gerner

## Abstract

Recently developed approaches for highly-multiplexed 2-dimensional (2D) and 3D imaging have revealed complex patterns of cellular positioning and cell-cell interactions with important roles in both cellular and tissue level physiology. However, robust and accessible tools to quantitatively study cellular patterning and tissue architecture are currently lacking. Here, we developed a spatial analysis toolbox, Histo-<u>Cyto</u>metric <u>M</u>ultidimensional <u>A</u>nalysis <u>P</u>ipeline (CytoMAP), which incorporates neural network based data clustering, positional correlation, dimensionality reduction, and 2D/3D region reconstruction to identify localized cellular networks and reveal fundamental features of tissue organization. We apply CytoMAP to study the microanatomy of innate immune subsets in murine lymph nodes (LNs) and reveal mutually exclusive segregation of migratory dendritic cells (DCs), regionalized compartmentalization of SIRPa^−^ dermal DCs, as well as preferential association of resident DCs with select LN vasculature. These studies provide new insights into the organization of myeloid cells in LNs, and demonstrate that CytoMAP is a comprehensive analytics toolbox for revealing fundamental features of tissue organization in quantitative imaging datasets.

## INTRODUCTION

Recent advances in intravital microscopy and multiplexed imaging approaches have revealed that the spatial organization of cell populations in tissues is highly complex and intimately involved in diverse physiological processes (e.g. cellular differentiation, organ function), as well as in major pathological conditions, such as infections, autoimmunity, and cancer. For the immune system in particular, cellular positioning is critical for both cell homeostasis and generation of protective responses during infection or after vaccination.^1–3^ Within lymph nodes (LNs) alone, different subsets of dendritic cells (DCs), professional antigen presenting cells, are spatially segregated within distinct tissue regions in a highly non-uniform fashion, which directly influences the sensitivity, kinetics, magnitude, and quality of the downstream adaptive immune response.^4–9^ Notably, advanced microscopy techniques have only recently revealed these findings in what were previously considered to be relatively well-studied organs, suggesting that further improvements in both microscopy and spatial analytics approaches can yield important insights into how complex biological systems operate.

This realization has inspired a number of emerging methods for highly multiplexed in situ cellular profiling (e.g. histo-cytometry, CODEX, CycIF, 4i, MIBI, STARMAP, spatial transcriptomics, etc).^4,10–20^ These techniques generate panoptic datasets describing phenotypic, transcriptional, functional, and morphologic cellular properties, while retaining information on the precise 2-dimensional (2D) or 3D positioning of cells within tissues. Analysis of images generated by these methods is complex and requires multiple processing steps, including image pre-processing, cell segmentation, object classification, and finally spatial analysis.^21–24^ Existing platforms excel at different steps of this pipeline, and much work is aimed at increasing the number and types of acquired analytes, enhancing image quality, as well as at improving cell segmentation to better define cellular boundaries.^21–25^ However, currently there is a lack of accessible and simple to use analytics tools for studying the complex multi-scale spatial relationships between different cell types and their microenvironments, for characterizing global features of tissue structure, as well as for understanding the heterogeneity of cellular patterning within and across samples. Existing approaches often utilize combinations of tools to reveal distance relationships between cells and tissue boundaries or utilize nearest neighbor and other statistical approaches to identify preferential interactions among different cell types.^10,11,15,22,26–28^ Together, the lack of readily accessible and easy to use analytics tools has hampered the ability of biologists with access to high-dimensional imaging technologies to obtain an in-depth understanding of the spatial relationships of cells and their surrounding tissue microenvironments within quantitative imaging datasets.

Here, we developed a user-friendly, spatial analysis method, Histo-<u>Cyto</u>metric <u>M</u>ultidimensional <u>A</u>nalysis <u>P</u>ipeline (CytoMAP), which utilizes diverse statistical approaches to extract and quantify information about cellular spatial positioning, cell-cell interactions and global tissue structure within imaging data. We implemented CytoMAP as a comprehensive toolbox in MATLAB, and it is specifically designed to analyze datasets generated with existing quantitative approaches, which already incorporate information on cell phenotype, morphology, and location. CytoMAP markedly simplifies spatial analysis by grouping cells into local neighborhoods, which can then be rapidly analyzed with diverse statistical approaches to reveal complex patterns of cellular composition, region structure and tissue heterogeneity. The CytoMAP platform incorporates multiple established and newly-developed statistical and visualization modules for analysis, including: machine learning based data clustering, cellular position correlation, distance analysis, tissue structure and heterogeneity visualization through dimensionality reduction, region network mapping, and 2D or 3D region reconstruction. Analysis with CytoMAP reveals and quantitates 2D or 3D tissue architecture, local cell composition and cell-cell spatial networks, as well as the interconnectedness of tissue regions. CytoMAP also facilitates robust sample-to-sample comparison, allowing exploration of structural and compositional heterogeneity across samples and diverse experimental conditions. Furthermore, CytoMAP can, in principle, be utilized for analysis of positionally-resolved data generated with diverse methods and across varying length scales, allowing integration into various disciplines.

We validate and demonstrate the capabilities and utility of CytoMAP by investigating innate and adaptive cell organization in steady state murine LNs, as well as in disease associated tissues, including solid tumors and *Mycobacterium tuberculosis* (Mtb) infected lung granulomas.^29–32^ Our analyses recapitulate previous descriptions of the cellular microenvironments within these tissues, as well as identify novel features of myeloid cell organization in LNs. Specifically, we reveal predominant localization of migratory SIRPa^−^ dermal DCs (dDCs) within the lower cortical ridge of LNs, as well as preferential association of LN-resident DCs with select LN blood vessels in distinct tissue regions.^33–36^ These findings uncover potential anatomical guidance cues that regulate myeloid cell localization, as well as suggest that LN vasculature may be heterogeneous based on the local interacting cell partners.^37,38^ Together, these data indicate that CytoMAP is a versatile and powerful tool for in-depth exploration of cellular positioning and tissue architecture.

## RESULTS

### CytoMAP enables analysis of tissue microenvironments, cellular interactions, and tissue structure

The conceptual basis for the spatial analysis used in CytoMAP centers on the notion that tissues are composed of cells that group together into local neighborhoods (Fig. 1a). These neighborhoods incorporate one or several cell types and groups of similar neighborhoods can extend across large distances to generate tissue regions. Unique combinations and interactions of regions further form the overarching tissue structure. Based on this concept, CytoMAP utilizes cell type and positioning information to phenotype neighborhoods and reveal how the distribution of cells and organization of neighborhoods leads to the generation of global tissue structure.

**Figure 1.**
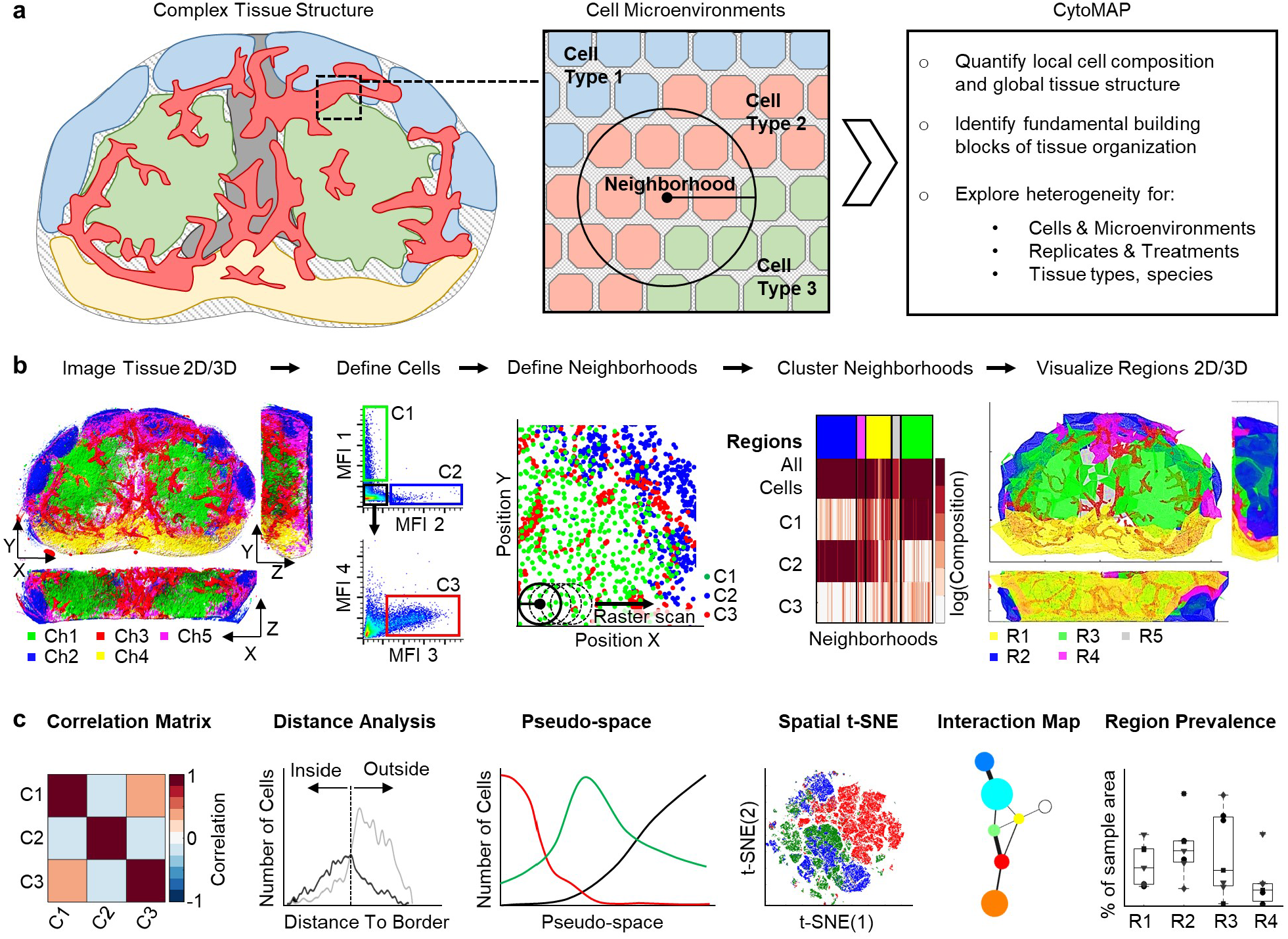
Workflow and features of CytoMAP. **a**, Conceptually, tissues are comprised of microenvironments defined by the local composition of different cell populations. CytoMAP is designed to extract quantitative information on cellular localization and composition within these regions, revealing how local cell microenvironments form global tissue structure, as well as allowing robust comparison of intra- and inter-sample tissue heterogeneity. **b**, The workflow starts with multi-parameter imaging of either thin sections or large 3D tissue volumes. Next, hierarchical gating of cell objects generated with existing pipelines is used to annotate distinct cell subsets, which are passed into CytoMAP for analysis. CytoMAP segments these spatial datasets into individual neighborhoods and uses clustering algorithms to define similar groups of neighborhoods, or tissue ‘regions’, which can be quantitatively explored and spatially reconstructed in 2D or 3D space. **c**, CytoMAP contains multiple tools to quantify and visualize the tissue architecture across length scales, ranging from analysis of spatial correlations between different cell types, investigation of distance relationships of cells with architectural landmarks, analysis of neighborhood heterogeneity within individual tissues or across multiple samples, as well as quantitative visualization of tissue architecture.

The basic CytoMAP workflow and currently incorporated analytical tools are presented in Fig. 1. For the tissues analyzed here, multi-parameter confocal microscopy and histo-cytometry were used to quantify the spatial location of immune cell subsets.^4,13,39^ The acquired phenotypic properties and positional information of individual cell objects are first passed to CytoMAP, which initiates the analysis process by spatially subdividing the cells into local neighborhoods of user-defined dimensions (Fig. 1b). The size of the neighborhood depends on the granularity of the desired information, with larger neighborhood size revealing more global patterns within the tissue, and smaller neighborhoods providing fine-grained information on hyperlocal cellular composition and tissue structure. If the dataset provided has no thickness (2D images), or thickness less than the user defined radius (thin section 3D data), CytoMAP utilizes a circular or cylindrical neighborhood window, respectively. A spherical neighborhood is used for analyzing larger volumetric 3D datasets. The generated neighborhoods contain information on the cell composition and density, expression of specific molecules, as well as data on any additional structural or functional parameters (e.g. local density of collagen fibers as detected by additional probes). These parameters are next passed to a self-organizing map (SOM) that clusters the neighborhoods into groups. This SOM clustering approach is neural network based, organizing information topologically and allowing extraction and quantification of critical features and unique tissue regions within highly complex datasets.^19,40–42^ The clustered neighborhoods represent areas within the tissue with similar cellular composition, and are thus defined here as tissue “regions”, which are denoted by the different colors in the top of the example heatmap in Fig. 1b. This heatmap allows direct visualization of the cellular composition of the neighborhoods (columns on the heatmap) in the identified regions, as well as shows the relative prevalence of the different regions within the imaged sample, as denoted by the size of the region color bars. Given that the individual neighborhoods also contain positional information (x,y,z) from the raster scan, they can be spatially remapped, as well as demarcated using the region color code. This allows direct visualization of the size and structure of the different regions within the tissue (Fig. 1b), and is different from conventional image display, as each region type is specified by a complex combination of multiple cell populations (defined in the heatmap), rather than by discrete imaged parameters.

A primary feature of CytoMAP is the incorporation of multiple visualization and quantification techniques, all encoded in a user-friendly interface in MATLAB, which collectively facilitate a more complete understanding of the spatial properties of cells, neighborhoods, and regions within tissues (Fig. 1c). In particular, local cell density within individual neighborhoods can be used to correlate the location of different cell types, revealing which cell populations preferentially associate with one another, or conversely avoid one another. This correlation analysis can be performed across multiple samples, and can be done either over entire tissues or within specified tissue regions. This is important, as cells may have distinct associations with one another in different tissue compartments. Distances of cells with respect to borders of tissue regions can also be easily evaluated (Fig. 1c), which provides information on the relative infiltration of cells into specific tissue compartments (e.g. T cell infiltration into tumors). Additionally, dimensionality reduction algorithms can be used to visualize tissue structure and complexity, as well as for sample-to-sample comparison. In particular, a new visualization approach, Pseudo-space, reduces the complexity of cell distribution across tissues into a one-dimensional plot, helping reveal the fundamental positional relationships of cells with respect to one another (Fig. 1c). Furthermore, spatial t-distributed Stochastic Neighbor Embedding (spatial t-SNE) analysis reduces the complexity of neighborhoods into a two-dimensional plot, facilitating comparison of the neighborhood heterogeneity within or across multiple samples (Fig. 1c).^43^ These dimensionality reduction techniques also help reveal how the tissue neighborhoods are organized to generate global tissue structure, which can be further quantified using region interaction network mapping (Fig. 1c) that calculates which regions preferentially border one another within the samples. Finally, CytoMAP allows comparison of region prevalence across different replicates or treatment conditions, which is a notoriously difficult task for heterogeneous samples with complex tissue architecture (Fig. 1c).

### CytoMAP quantifies well-defined tissue structure in LNs

We first validated the CytoMAP workflow by analyzing murine LN tissues, which have well-defined organization of basic immune cell types.^2^ To accomplish this, a 20μm thick section of a draining LN from a C57BL/6 immunized mouse was stained with a panel of directly conjugated antibodies against distinct innate and adaptive immune populations and imaged using a confocal microscope. The image in Fig. 2a shows staining of the tissue with markers for B cells (B220), DCs (CD11c), and T cells (CD3), demonstrating pronounced segregation of these cell types to previously defined tissue compartments.^2^ Individual cells within the image were next segmented in 3D, and the mean fluorescent intensity (MFI), as well as the (x,y,z) positioning of the resulting cell objects were imported into FlowJo for hierarchical gating of three primary cell types: T cells, B cells, and CD11c-expressing cells, which are mainly DCs (Fig. 2b), as described previously.^4,13,39^ Next, the positional data on these cell populations were imported into CytoMAP for further processing. In CytoMAP the cells were subdivided into 30μm radius neighborhoods using the *Raster Scan Neighborhoods* function, which digitally raster scanned a cylindrical window with the user-defined radius over the dataset (Fig. 2c). This neighborhood radius size was chosen empirically, as it provided an optimal balance of spatial granularity and processing speed to reveal fundamental features of cellular organization for this specific sample, and was consistent with biological data on the dispersion distances of secreted cytokines.^44^ A SOM was next used to cluster these neighborhoods based on the cellular composition (number of cells of each type in a neighborhood divided by the total number of cells in that neighborhood), but not utilizing the position of the neighborhoods within tissues. Exclusion of neighborhood positions is important for comparison of compositionally similar, but spatially distal neighborhood types. Given that the number of regions used for SOM clustering requires user input and thus can be subjected to bias, we incorporated statistical tools to determine the optimal number of regions needed to reveal the underlying data structure. In this example, the number of regions was determined using the Davies-Bouldin criterion, which uses the ratio of within-cluster to between-cluster distances.^45,46^ The heatmap in Fig. 2d shows the cell composition (rows) of the individual neighborhoods (columns) and which cluster/region they were assigned to (top color bar). This analysis identified different tissue regions that were primarily composed of either B cells, T cells, DCs, or those with mixed cellular composition. These regions were next visualized by plotting the positions of the color-coded neighborhoods (Fig. 2e). This plot demonstrates reconstruction of the original image, revealing localization of computationally defined B cell follicles (blue), deep T cell zone (red), the outer T zone paracortex and the T-B border (orange), as well as the LN medullary and subcapsular regions (green). Of note, the red/orange regions at the top of the reconstructed image (Fig. 2e) correspond to noise in the original confocal image (Fig. S1a). In computationally heavy image processing pipelines, special care must be taken to either accurately remove such artifacts or to ensure appropriate interpretation of the results. Together, this indicates that CytoMAP can be used for unbiased identification of different tissue regions in a scalable and reproducible manner, eliminating the need for manual image annotation and allowing quantitative evaluation of larger sample sets.

**Figure 2.**
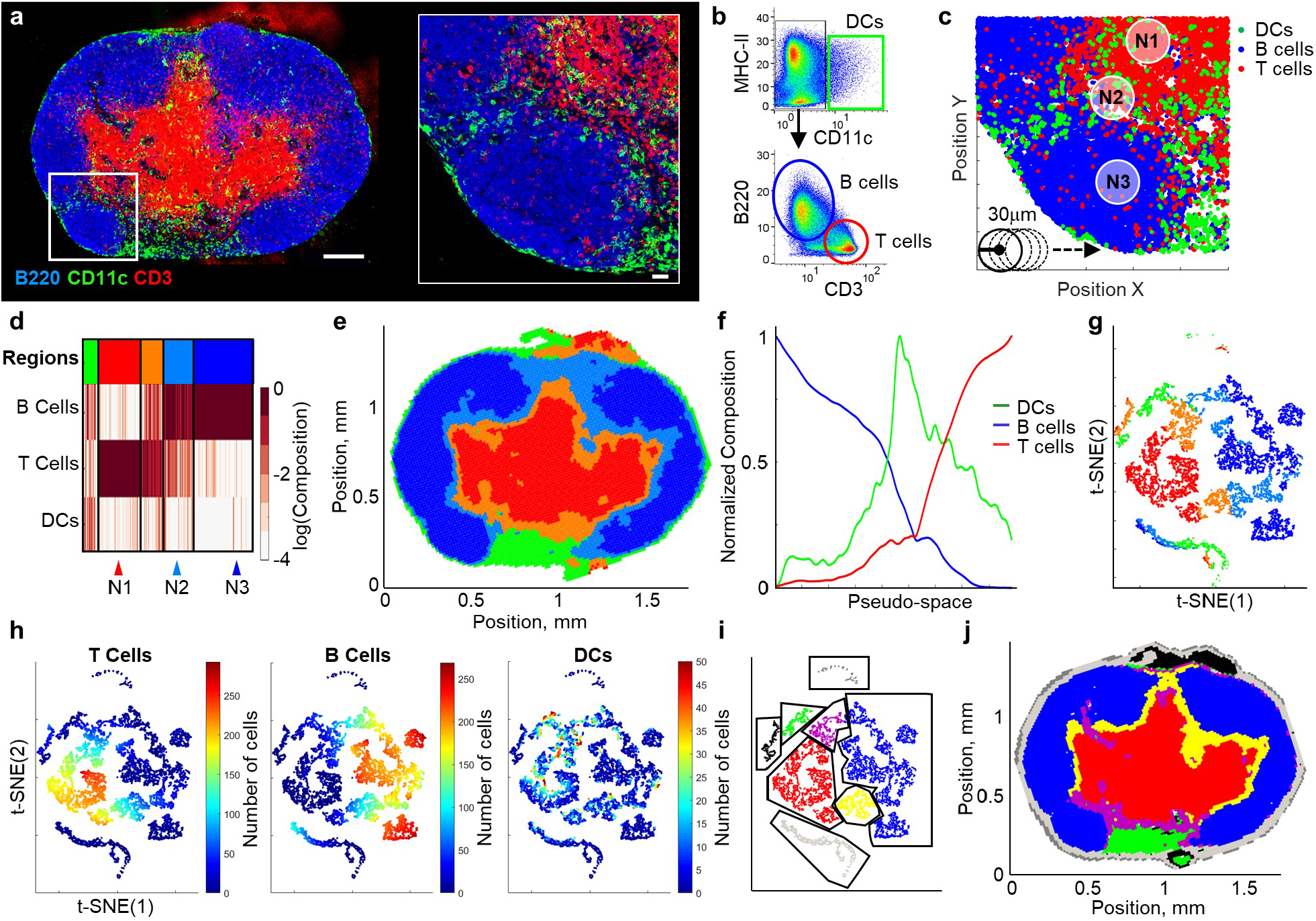
CytoMAP automatically identifies major features of LN tissue structure. **a**, Multi-parameter confocal microscopy image, and zoom-in, of a LN section from an immunized C57BL/6 mouse demonstrating highly compartmentalized staining for B220 (B cells), CD11c (DCs) and CD3 (T cells). Overview image scale bar = 200μm; zoom-in scale bar = 30μm. **b**, Cell objects within the image were segmented and analyzed using histo-cytometry. Plots demonstrate gating used for identification of the indicated immune cell populations. Here, cells with high CD11c MFI are referred to as DCs, albeit this gate may include additional CD11c-expressing cell types. **c**, Cell data from panel b were passed into CytoMAP and positionally plotted (shown area matches panel a zoom-in). CytoMAP was used to raster scan the cells into 30μm radius neighborhoods based on their spatial position, as denoted by the arrow. **d**, Heatmap of the neighborhood cellular composition (percentage of each cell phenotype per neighborhood) after SOM clustering. Individual clusters, ‘regions’, are denoted by the color bar at the top of the graph. Arrowheads at the bottom highlight the specific neighborhoods also shown in panel c. **e**, Region color-coded positional plot of the neighborhoods from panel d demonstrating general reconstruction of LN architecture. **f**, Pseudo-space plot with the neighborhoods sorted based on B cell composition (sorted to the left) and T cell composition (sorted to the right) on the Pseudo-space axis. This plot demonstrates an abstracted representation of cellular distribution across different LN compartments. **g**, t-SNE plot of the neighborhoods, in which the numbers of cells and total MFI of all channels, but not positional information, were used for the dimensionality reduction. Neighborhoods were color-coded based on region type, as identified in panel d. **h**, Same t-SNE plot as in panel g, but with the neighborhoods color-coded as a heatmap reflecting the number of the indicated cell types per neighborhood. **i**, The t-SNE plot was manually gated based on the cell composition visualized in panel h. **j**, Gated neighborhoods were positionally remapped using the color code definitions from panel i. For this experiment, an imaging volume of 0.03 mm3, 139,399 cells, and 11328 neighborhoods were analyzed.

In addition to discreet region definitions, we implemented a data visualization function, *Pseudo-space*, to visualize the continuous region transitions within tissues, and investigate how cells are distributed across different tissue compartments (Fig. 2f). Pseudo-space allows the user to sort the neighborhoods by the absolute number or composition of different cell types within the neighborhoods and plots the neighborhoods in this sorted order on a linear Pseudo-space axis. This provides a qualitative picture of how different cell types change in their composition across the neighborhoods along this user defined dimension. For the LN analysis, the neighborhoods were sorted such that B cell rich neighborhoods fell to the left and T cell rich neighborhoods fell to the right along the Pseudo-space axis. Cellular composition of the neighborhoods was then normalized and smoothed along the Pseudo-space axis, aiding the visualization of cell populations with different tissue abundance and with substantial heterogeneity across the tissue. Pseudo-space visualization demonstrated that as the percentage of B cells in the neighborhoods declined, the percentage of T cells increased (Fig. 2f), mirroring what was observed in the original image (Fig. 2a). In the transitional area between the B cell and T cell rich neighborhoods, Pseudo-space analysis revealed an increased portion of DCs, which likely corresponds to the increased abundance of DCs in the medullary and paracortical regions of the LN. Thus, Pseudo-space provides the user with a way to reduce the complexity of tissue structure to a single dimension, aiding in visualization of cellular relationships within complex tissues and linearizing complex microenvironments.

In addition to manual dimensionality reduction, we implemented t-SNE dimensionality reduction to explore the neighborhood heterogeneity based on cellular composition and biomarker expression on a two-dimensional plot (Figs. 2g, 2h, S2b). Instead of individual cells, as is typically done, the cellular composition of the neighborhoods, but not their position, was used for the t-SNE analysis. This revealed complex structure in the LN dataset with clearly demarcated, but also interconnected clusters of neighborhoods within the t-SNE 2D space (Fig. 2g). Color-coding of neighborhoods based on region clustering, as defined in Fig. 2d, revealed clear association of the distinct regions with the different clusters generated by the t-SNE analysis, suggesting that both methods are capable of identifying cellular organization within tissues. This was confirmed using manual gating, directly within the t-SNE plot (Fig. 2i), and spatial mapping (Fig. 2j) of the t-SNE clusters, which accurately reconstructed the global tissue architecture, as well as clearly identified image artifacts (black color-coded gate Fig. 2i-j). t-SNE analysis also demonstrated the inter-connected nature of the regions, with the neighborhood regions assigned to the B cell follicle and T cell zone regions being connected with one another by neighborhoods assigned to the paracortical region (orange, Fig. 2g). This is likely due, in part, to smoothing effects from raster scanning the neighborhoods in steps of half the defined radius, such that neighborhoods partially overlap. However, this t-SNE data structure also captures features of the actual tissue organization, identifying the paracortical T-B border regions where T cells and B cells are in sufficient spatial proximity to be included in the same neighborhood (Fig. 2c, neighborhood N2). It is likely that lowering the radius scan size would reduce but not eliminate the interconnectedness of these regions, as even in tissues with extremely sharp compartment boundaries, individual cells would still exhibit differential neighboring partners at the border vs. the center of that compartment. Thus, the t-SNE analysis is able to reveal features of tissue organization based on neighborhood cell composition without being provided information on the positioning of that neighborhood within the sample.

We next implemented network analysis to interrogate the interconnectedness of the different regions with one another by calculating the percentage of the region borders that are shared with other regions (Fig. S1c). Mapping of these bordering relationships revealed that the T cell zone regions (red) were connected to the B cell regions (blue) via the paracortical regions (orange), thus directly recapitulating the t-SNE analysis and the original image (Fig. S1c, 2a, 2g). Together, these proof-of-concept data indicate that the CytoMAP analysis tools were capable of robustly identifying the key features of cellular organization and tissue structure for relatively simple cell types with well-characterized spatial properties.

### CytoMAP analysis of the tumor microenvironment

We next tested the capabilities of CytoMAP in exploring the distribution of immune cells in more complex tissue types. For this, we imaged a cross section of a whole murine CT26 colorectal tumor stained with a panel of markers to detect various innate and adaptive immune populations (Fig. 3a). Histo-cytometry was used to gate on T effector cells (Teff), T regulatory cells (Treg), B cells, tumor associated macrophages (TAMs), activated macrophages (aMacs), DCs, and MHC-II^+^ SIRPa^DIM^ myeloid cells (Fig. S2a). The spatial distribution of the cell populations revealed striking compartmentalization of the tumor tissue into areas enriched with different immune cell subsets (Fig. 3b). SOM clustering in CytoMAP of 50μm raster scanned neighborhoods identified tissue regions preferentially associated with specific myeloid cell populations (Fig. 3c, S2b). Spatial visualization of these regions further revealed discrete tumor zones composed of relatively segregated region types associated with distinct cell subsets (Fig. 3d). As an example, region R6 was predominantly composed of DCs and Teff cells, and was primarily localized to the outer periphery of the tumor (Fig. 3c, 3d), resembling the lymphocytic cuff previously observed in colorectal and other cancers.^30^ In contrast, regions R2 and R3 were dominantly composed of TAMs, and these regions were localized within the deeper portions of the tumor (Fig. 3c, 3d). To better understand the spatial relationships among these tumor microenvironments, we next utilized the *Pseudo-space* function. For this, smoothed distributions of neighborhoods enriched in DCs were sorted to the left and those enriched in TAMs were sorted to the right along the Pseudo-space axis. This analysis revealed the underlying distribution of the different immune subsets across the tissue, demonstrating increased abundance of TAMs in the neighborhoods more proximal to the tumor core, and with increased presence of lymphocytes and DCs in the tumor periphery (Fig. 3e). Interestingly, Teff cells were well-represented in both the peripheral immune cuff and the tumor core, while the Tregs were predominantly restricted to the outer tumor periphery. This indicates that this CT26 sample represents a relatively well-infiltrated ‘hot’ tumor that may be susceptible to checkpoint blockade therapy, which is consistent with published observations.^47,48^ These spatial relationships were further quantified using the Pearson correlation coefficients of the number of cells per neighborhood (Fig. 3f, S2c). This demonstrated positive correlation of T and B cells with DCs and aMacs, and negative correlation with TAMs, thus revealing the local cellular networks and the preferential associations of different cell types across distinct intra-tumor microenvironments. Notably, the negative correlation of Teff cells and TAMs indicated that while Teff cells were generally capable of infiltrating the tumor tissue, they were not enriched in the areas dominantly populated with TAMs, and this was also seen in the Pseudo-Space plot, in which the Teff numbers dropped in the TAM-associated neighborhoods (Fig. 3e). This indicates that while generally capable of partial tumor infiltration, Teff lack the ability to infiltrate the deep tumor nests in this cancer model. Collectively, these data reveal marked segregation of different myeloid cell types across the tumor tissue, as well as demonstrate that the different analytical modules in CytoMAP can be successfully used to study cellular localization within complex tissues.

**Figure 3.**
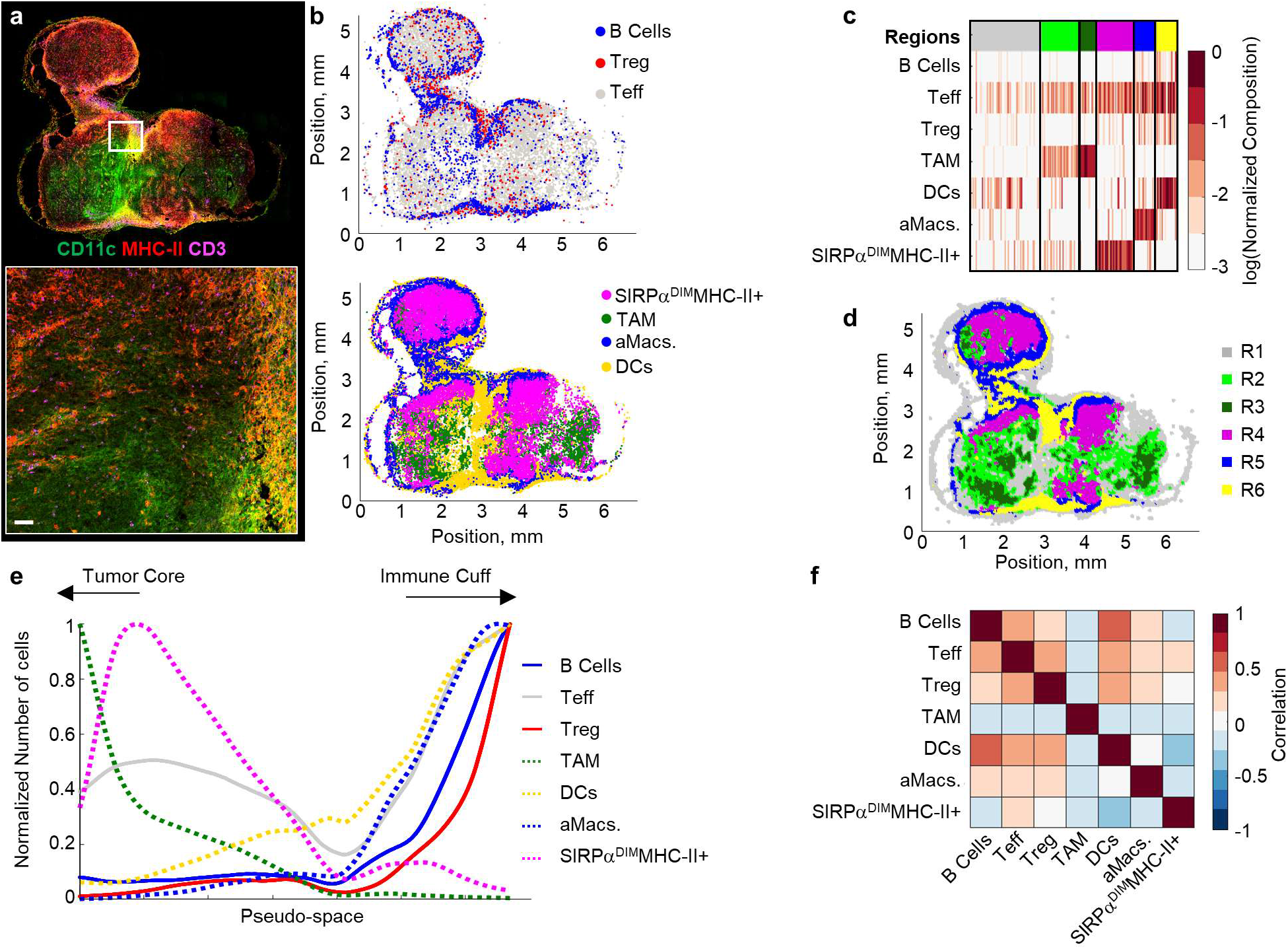
CytoMAP analysis of a murine colorectal tumor sample. **a**, Multiplex confocal image of a 20 μm thick CT26 tumor section isolated 9 days after subcutaneous inoculation and stained with the indicated markers. Zoom in demonstrates the immune cell cuff surrounding the tumor. Main image scale bar = 500μm; zoom-in scale bar = 50μm. **b**, Positional plot of the lymphocyte (top) and myeloid cell (bottom) populations as defined by histo-cytometry gating presented in Fig. S2a. **c**, Heatmap of the normalized immune cell composition of regions defined by SOM clustering of 50μm radius neighborhoods. **d**, Positional plot visualizing the CT26 color-coded regions defined in panel c. **e**, Pseudo-space plot with the number of the indicated cell types per neighborhood after pre-sorting for TAM (sorted to the left) and DCs (sorted to the right) along the pseudo-space linear axis. Neighborhoods were also smoothed, and yaxis normalized to allow qualitative comparison of cell type localization. **f**, Heatmap of the Pearson correlation coefficients of the number of cells per neighborhood across the imaged tumor sample. For this experiment, an imaging volume of 0.3 mm^3^, 34,013 myeloid cells, 22,035 lymphocytes, and 58,608 neighborhoods were analyzed.

### CytoMAP reveals underlying organization of immune cells in Mtb granulomas

In addition to tumors, previous studies have demonstrated structured organization of immune cells within Mtb-infected pulmonary granulomas.^29,31,49–51^ We thus tested the ability of CytoMAP to quantitatively explore such organization in a 20μm lung section from a mouse infected with aerosolized Mtb. We observed formation of a discrete Mtb-associated lung granuloma and complex patterns of cellular distribution within this structure (Fig. 4a). Two regions of interest, one of the unaffected lung and one of the Mtb-infected granuloma (Fig. 4a), were imaged at high resolution for in depth analysis. As before, histo-cytometry was used to gate on the different cell populations, including CD4^+^ and CD4^−^ T cells, B cells, CD11b^+^ myeloid cells, alveolar macrophages (Alv. Macs.), DCs, and Mtb^+^ infected cells (Fig. S3a). The positions of these cells were next passed into CytoMAP for remapping (Fig. 4b) and analysis. A small number of the Mtb^+^ objects were identified outside the granuloma in the images. These appeared extra-cellular and likely represented imaging artifacts; they were not selectively removed from analysis to avoid introducing user bias. Heatmap visualization of the clustered, 50μm radius raster scanned neighborhoods, with automatic region number identification, revealed distinct region types composed of different immune cell populations (Fig. 4c, S3b, S3c). Remapping of the region color-coded neighborhoods demonstrated that the neighborhoods enriched in Mtb-infected myeloid cells (R3) were primarily located in the deep center of the granuloma (Fig. 4d), recapitulating visual observations. These infected regions were surrounded by neighborhoods containing high concentrations of T cells (R4 and R5), and these were further surrounded by regions associated with uninfected myeloid cells and Alv. Macs. (R2). We also detected segregated B cell rich neighborhoods in region R6 (Fig. 4d), which were reminiscent of B follicles and tertiary lymphoid structures. To further visualize these cellular relationships, we performed Pseudo-space analysis (Fig. 4e). This plot demonstrated that within the granuloma, T cell and B cell rich neighborhoods were concentrated just outside of the core infected Mtb^+^ cells. The Alv. Macs and DC-rich neighborhoods also appeared excluded from the central core of the granuloma, recapitulating the image data. These relationships were also quantitatively explored by calculating the Pearson correlation coefficients of the numbers of cells in the neighborhoods (Fig. 4f, S3d). In this analysis, the left half of the heatmap, representing uninvolved lung tissue, demonstrated no strong correlations between the different immune cell types, consistent with qualitative observations (Fig. 4a, 4b). In contrast, the right half of the heatmap, representing the granuloma, demonstrated positive correlation between the Mtb^+^ infected cells and the CD11b^+^ myeloid cells, weaker correlation with the surrounding T cells, and negative correlation with Alv. Macs. These findings are consistent with previous observations of immune cell organization in Mtb granulomas, describing the partial segregation of CD4^+^ T cells from Mtb-infected cells and the formation of tertiary lymphoid structures.^49^ Collectively, these data indicate that CytoMAP is capable of robust analysis of highly complex tissue structures across diverse organ types and disease settings.

**Figure 4.**
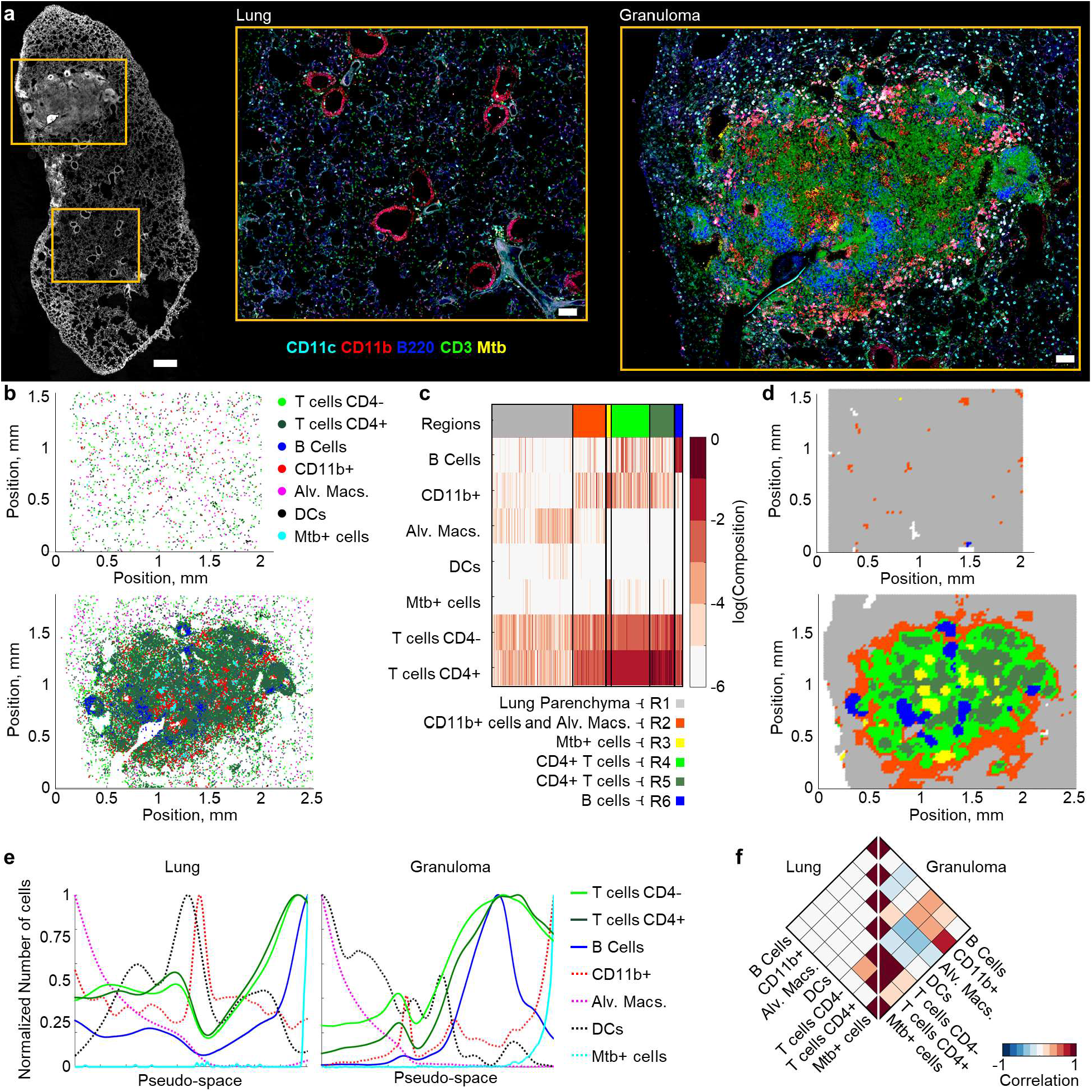
CytoMAP analysis of Mtb-infected lung granuloma. **a**, Multiplex confocal image of a 20μm thick section from a Mtb-infected murine lung sample stained with the indicated antibodies. The left image shows multiple imaged channels overlaid in white to visualize the overall lung structure. Scale bar = 500μm. Zoom in images demonstrate separately acquired regions of interest centered on a representative uninvolved lung area and the Mtb granuloma. Zoom-in scale bar = 100μm. **b**, Positional plots of the immune cell subsets in the uninvolved (top) versus granuloma-containing (bottom) lung areas shown in panel a, with the cell populations defined by the histo-cytometry gating scheme presented in Fig. S3a. **c**, Heatmap of SOM-clustered, 50μm radius neighborhoods demonstrating the distinction between uninvolved lung regions (R1) versus those within the granuloma (R2-R6). **d**, Positional plots of the neighborhoods, color-coded by region defined in panel c, within the uninvolved (top) versus granuloma-containing (bottom) lung areas. **e**, Pseudo-space plots visualizing the number of the indicated cell types per neighborhood within the uninvolved (left) versus granuloma-containing (right) lung areas after sorting for Alv. Macs (sorted to the left) and Mtb^+^ cells (sorted to the right) along the pseudo-space axis. Neighborhoods were also smoothed, and y-axis normalized to allow qualitative comparison of cellular associations. **f**, Rotated half-heatmaps demonstrating the Pearson correlation coefficient of the number of cells within the neighborhoods across either the uninvolved lung (left) or the granuloma (right). For the uninvolved lung region, an imaging volume of 0.05 mm^3^, 36,194 cells, and 4725 neighborhoods were analyzed. For the granuloma lung region, an imaging volume of 0.07 mm^3^, 140,453 cells, and 7,350 neighborhoods were analyzed.

### Myeloid cell organization in steady state LNs

Finally, to test the performance of CytoMAP on larger datasets with complex cellular organization, we turned to investigating the localization of different myeloid cell subsets in a cohort of steady state LNs, as these cell populations have been previously shown to display intricate, non-overlapping distribution patterns within these tissues.^2–5^ To this end, 20μm sections from steady state LNs were stained with a 12-plex antibody panel to detect distinct DC and macrophage subsets, as well as to visualize T cells, B cells, and different blood and lymphatic stromal cells as reference structural markers. As above, tissues were imaged (Fig. 5a) and distinct cell populations were identified by histo-cytometry (Fig. 5b, S4a). Positional and MFI data on these cell types were next passed to CytoMAP for analysis. Basic spatial remapping of these populations in CytoMAP recapitulated the general LN structure, as well as qualitatively validated previous findings on the location of different DC and macrophage subsets (Fig. 5c). In particular, we observed that subcapsular sinus (SCS) and medullary (Med) macrophages localized to the outer LN periphery and medullary regions, respectively. Resident cDC1 and cDC2 populations also exhibited previously established spatial patterns, with the resident cDC2s preferentially localizing in peripheral LN regions, and the resident cDC1s more evenly distributed across the LN parenchyma and within the T cell zone (Fig. 5c, S4b).^4,7,52^ Similarly, visualization of the migratory DC subsets confirmed previous observations, with CD207^+^ cells (Langerhans cells and migratory cDC1s) located within the central T cell zone, and the CD301b^+^ and SIRPa^+^ dermal DC (dDC) subsets located in regions bordering the B cell follicles and in the lower cortical ridge, respectively (Fig. 5c, S4b).^4,6,7,53,54^ In addition, histo-cytometry analysis revealed a recently described population of migratory SIRPa^−^ dDCs, although the spatial positioning of this population has not been established in these previous studies.^33,34^ Qualitative visualization of this migratory DC subset revealed that these cells were predominantly localized in the lower cortical ridge bordering the LN medulla (Fig. 5c, S4b). Visual inspection of the original confocal data confirmed abundant presence of SIRPa^−^ migratory dDCs in this LN compartment, with these cells forming a dense cuff-like cellular aggregate at the border of the T cell zone and the LN medulla (Fig. 5a).

**Figure 5.**
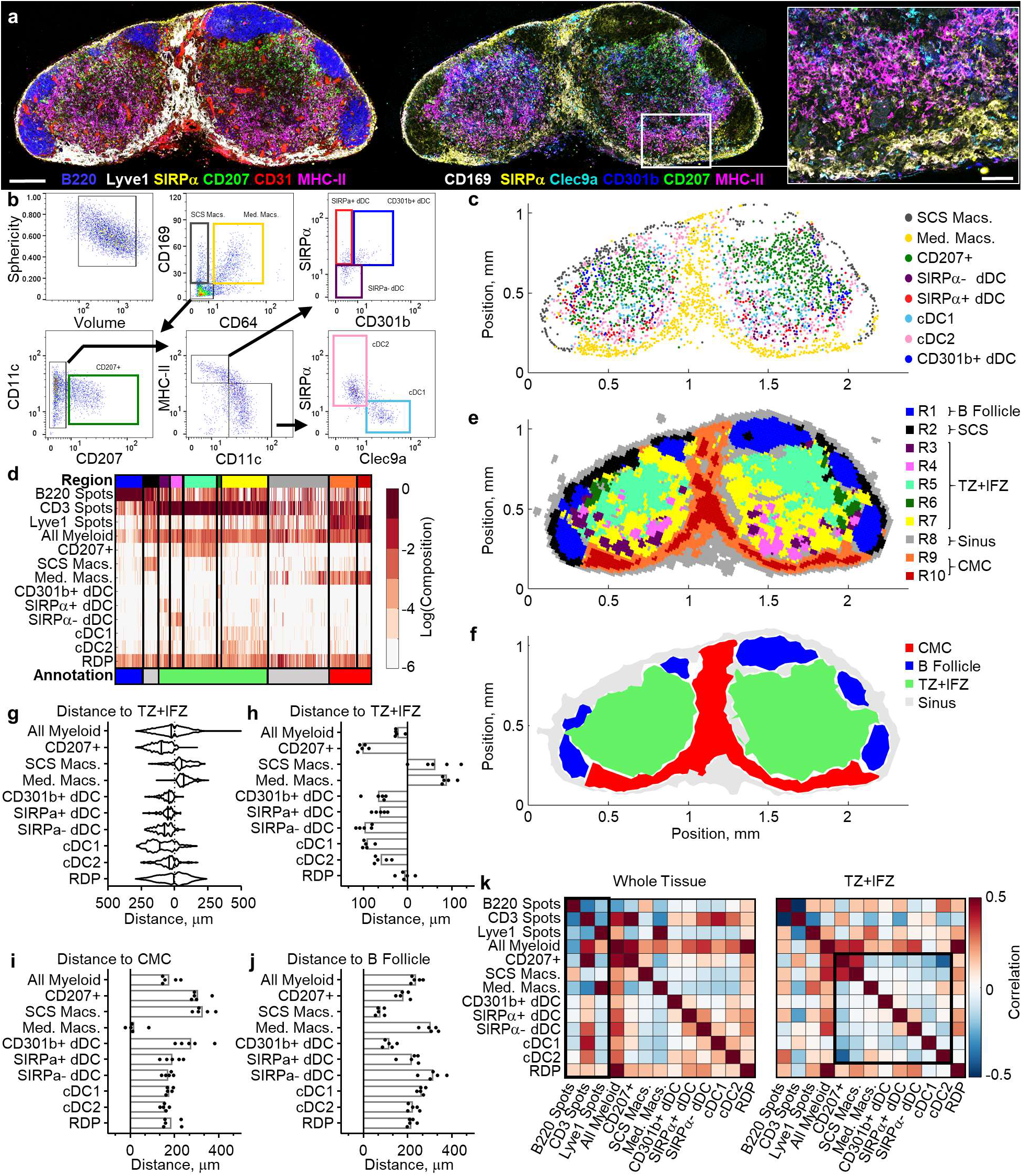
CytoMAP reveals diverse patterns of myeloid cell organization in LNs. **a**, Multi-parameter confocal microscopy image of a representative steady state non-draining LN section from an immunized C57BL/6 mouse. Images demonstrate staining for the indicated myeloid cell markers, with the image on the left also showing B220 (B cells), Lyve1 (lymphatics) and CD31 (blood vessels) staining. Scale bar = 200μm. The right zoom-in demonstrates a cellular aggregate of SIRPa^−^ dDC in the lower T cell zone paracortex proximal to the LN medulla. Scale bar = 50μm. **b**, Histo-cytometry gating scheme used to annotate the myeloid cell subsets within the imaging data. **c**, Cellular positions plotted in CytoMAP showing the distinct localization patterns of the indicated immune cell subsets. **d**, The heatmap of SOM clustered, 30μm radius neighborhoods demonstrates different LN regions (top color bar) as defined by the distinct composition of myeloid cells. Regions were also manually annotated (bottom color bar) reflecting the association with B220, CD3 or Lyve1 landmark spots. Annotated gating shown in Fig. S5c. **e**, Positional plot of the LN neighborhoods as color-coded by the region type (panel d - top color bar). **f**, Positional plot of surfaces generated on the manually annotated regions (panel d - bottom color bar). **g**, Violin plot showing the number of cells as a function of distance to the border of the TZ+IFZ annotated region for the representative LN sample. Distances to the left of zero represent cells inside the region; distances to the right of zero represent cells outside the region. **h**, Plot of the distances of cells to the TZ+IFZ region, in which each dot represents the mean distance of the indicated cell population in an individual LN sample (n=5). **i**, Mean distances of indicated cells to the CMC region. **j**, Mean distances of indicated cells to the B Follicle region. **k**, Correlation plot demonstrating the Pearson correlation coefficients between the number of indicated cells per neighborhood using either all tissue neighborhoods (left) or only the TZ+IFZ region neighborhoods (right). Correlations were averaged over the sample cohort (n=5) from one experiment. For this experiment, an average imaging volume of 0.023 mm^3^, and an average of 6,611 myeloid cells, 84,347 spots, and 12,708 neighborhoods were analyzed per sample. Data are representative of at least two independent experiments.

CytoMAP was next used to quantify features of cellular organization within these steady state LNs. To aid in accurate classification of the tissue regions, additional spot objects were generated based on different landmark channels demarcating known LN structures (i.e. CD3, B220, and Lyve1 for the T cell zone, B cell follicles and lymphatic vessels, respectively), and the positional data from these landmark objects were also passed into CytoMAP. In addition, using the *Generate Random Points* function in CytoMAP, a population of randomly distributed points (RDP) were defined throughout the LN and were used to compare to the distribution of the imaged cells (Fig. S4b). As above, the cells and landmark spots were subdivided into individual neighborhoods by digitally raster scanning a 30μm radius neighborhood filter using the *Raster Scan Neighborhoods* function. These neighborhoods were next clustered using a SOM, with the number of regions automatically determined by the Davies-Bouldin criterion. This resulted in generation of 10 distinct regions (color bar at the top of Fig. 5d) that were enriched with different cell types, suggesting a relatively discrete distribution of cells within the LNs. Positional remapping of the neighborhoods, color-coded by region, directly recapitulated the spatial boundaries and the general architectural features seen qualitatively in the original image, suggesting robust digital tissue reconstruction (Fig. 5e, S5a, S5b). This analysis also demonstrated that the different DC and macrophages subsets appeared relatively segregated from one another within distinct spatial compartments, corroborating previous observations.^4^ Region visualization also established the predominant localization of the SIRPa^−^ migratory dDC population (R4 region) in the lower cortical ridge bordering the LN medulla (Fig. 5e, S5a, S4b).

To further delineate the distribution of different cell types across physiologically defined LN compartments, we used CytoMAP to manually annotate the regions based on landmark spot clustering (Fig. 5d, S4b, S5c). This resulted in new composite regions, color annotated at the bottom of the heatmap in Fig. 5d, with region R1 corresponding to B cell follicles, region R2 to the sub-capsular sinus (SCS), regions R3-R7 to the T cell zone and Interfollicular zones (TZ+IFZ), region R8 to the sinus, and regions R9 and R10 to the cortico-medullary cords (CMC) (Fig. 5d, 5e, S5c). Next, using the *Make Surface* function in CytoMAP, we built surfaces around the neighborhoods belonging to these annotated groups (Fig. 5f), then calculated the distances of the myeloid cells to the borders of these surfaces (Fig. 5g, S5e). The distance to each region border for all cells in a single sample (Fig. 5g, S5e) or averaged over all cells for multiple samples (Fig. 5h-j, S5f), corroborated the qualitative observation that most DCs were distributed within the TZ+IFZ, while the macrophages were positioned in either the SCS or CMC. This analysis also confirmed preferential localization of resident cDC2s in closer proximity to the CMC and B cell follicles, compared to the more heterogeneous distribution of resident cDC1s within the LN (Fig. 5i, 5j). Additionally, distance analysis revealed that CD301b^+^ DCs were primarily distributed in close proximity to the B cell follicles, as previously described,^53^ while the SIRPa^−^ migratory dDCs were located distally from the B follicles and in close proximity to the CMC (Fig. 5i, 5j, S5f).

The discrete clustering and segregation of different myeloid cell types (Fig. 5d, 5e) also indicated that the DC subsets were distributed in spatially non-overlapping patterns. To explore this further we calculated the Pearson correlation coefficients of the number of cells per neighborhood, averaged over all of the samples, for either the whole samples or only the neighborhoods within the TZ+IFZ regions (Fig. 5k, S5g). Whole tissue correlation analysis demonstrated that all DC populations were positively correlated with CD3 spots and negatively correlated with Lyve1 spots and medullary macrophages (Macs), indicating that on average, most DCs are positioned in the T cell zone and not in the LN medulla (Fig. 5k left). SCS Macs were also positively correlated with B220 spots, in line with their enhanced prevalence lining the B cell follicles. Correlation analysis of the TZ+IFZ compartment, enriched in most DC populations, revealed that migratory DC populations (i.e. CD207^+^ DCs, CD301b^+^ dDCs, SIRPa^+/−^ dDCs) were in general negatively correlated with one another or displayed little spatial correlation (Fig. 5d, 5k right). This provides quantitative support to the observation of spatial segregation of DC subsets in LNs. Together, these analyses highlight the ability of CytoMAP to delineate complex patterns of cellular organization into quantitative metrics. Biologically, these observations reveal strong spatial exclusivity for different DC populations, as well as identify the distribution profile of migratory SIRPa^−^ dDCs.

### Spatially organized myeloid cell associations with LN blood vessels

In addition, qualitative examination of imaged LN sections revealed that some DC populations appeared to be highly proximal to LN blood vessels. Given the established role of DCs in homeostatic maintenance of LN vasculature, we next examined the relative distribution of DC subsets with respect to LN blood vessels.^35,36^ To account for the sampling error associated with thin section imaging, in which critical tissue landmarks may lie just above or below the sectioning plane, we turned to volumetric microscopy of stained and Ce3D optically-cleared 500 μm thick slices of steady state murine LNs.^13,39^ This resulted in robust visualization of complex patterns of DC subsets in 3D tissue space (Fig. 6a). Qualitative examination of the imaged LNs revealed close association of the Clec9a^+^ resident cDC1s with CD31^+^ vascular endothelial cells, and many of these DCs appeared to encapsulate large segments of the neighboring blood vessels (Fig. 6a and Video S1). In contrast, CD207^+^ migratory DCs (Langerhans cells and migratory cDC1s) appeared less associated with the LN vasculature. To quantitate these observations, we performed histo-cytometry to identify various DC and macrophage populations and passed the positional data of these cells to CytoMAP (Fig. S6a, S6b).^13,39^ To provide positional information on CD31^+^ blood vessels, we generated segmented surface objects on the CD31 channel and imported these objects’ data into CytoMAP. As above, positional information of B220^+^ B cell and Lyve1^+^ lymphatic sinus landmark spots were also included in the analysis. In addition, RDPs were defined throughout the 3D LNs for comparison. The distances for the different myeloid cell subsets, or RDPs, to the nearest blood vessel object were next calculated in CytoMAP (Fig. 6b). Visualization of the individual cell distances to the closest blood vessel demonstrated that, while there was substantial heterogeneity within a given cell population, both resident cDC1 and cDC2 populations were on average located in close proximity to blood vessels (Fig. 6a-d). This was in contrast to more distal relationships for most migratory CD207^+^ DCs, Macs, or RDPs. To further explore these relationships, we calculated the Pearson correlation coefficients for the number of cells per 50μm radius neighborhood. This analysis demonstrated that in contrast to migratory DCs, resident cDC1 and cDC2 subsets were both positively correlated with blood vessels (Fig. 6e, S6c). These data indicate that resident DCs are spatially associated with LN vasculature, suggesting that blood vessels may provide guidance cues to guide the localization of these myeloid cell types in LNs.

**Figure 6.**
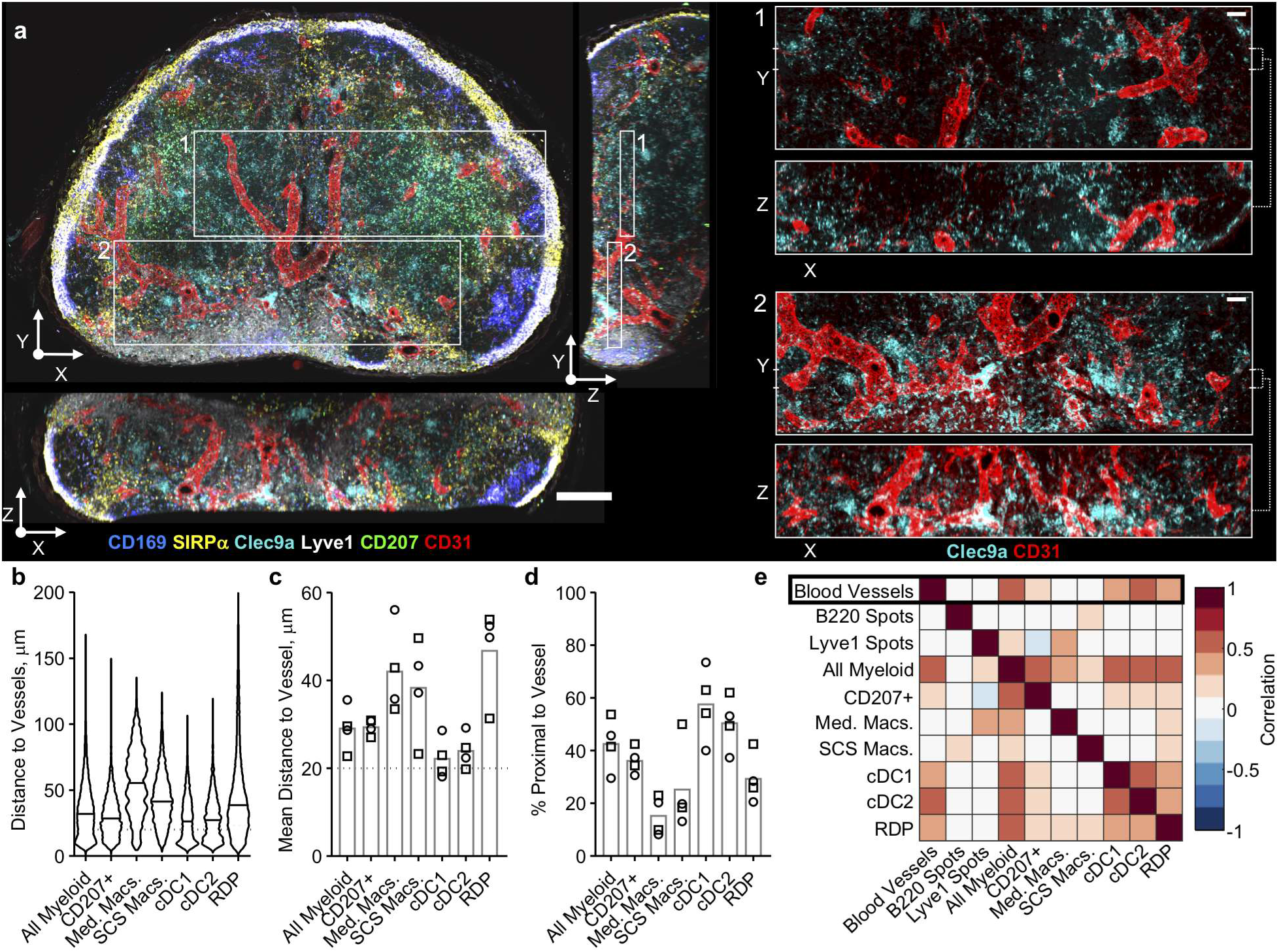
CytoMAP analysis of 3D LNs reveals preferential interactions of DCs with vasculature. **a**, Representative confocal image of a 500μm thick, Ce3D cleared steady state LN slice. Zoom-in images demonstrate association of Clec9a^+^ cDC1 cells with CD31^+^ blood vessels. Main image scale bar = 200μm; zoom-in scale bar = 50μm. **b**, Violin plot comparing the distances of the indicated cells to nearest blood vessels within a representative LN sample. **c**, Mean distances of cell populations to the nearest blood vessel, with each symbol representing an individual LN sample. Circles and squares represent samples from two independent experiments. Cells below the dotted line at 20μm were considered proximal to vasculature. **d**, Percentage of proximal cells for the LN samples shown in c. **e**, Heatmap showing the Pearson correlation coefficients, averaged across all imaged samples, between the number of different cell or landmark object types per 50μm radius neighborhood. Data represent four samples from two independent experiments. For these experiments, an average imaging volume of 0.93 mm3, and an average of 39,139 myeloid cells, 36,048 blood vessel objects, 98,279 spots, and 133,446 neighborhoods were analyzed per sample.

The spatial segregation of resident cDC1 and cDC2 subsets into distinct regions of the LN (Fig. 5c, S5b) also suggested that blood vessels may be differentially associated with specific DC populations across the distinct tissue compartments. To explore the cellular microenvironments associated with the complex vascular networks in LNs (Fig. 7a), we utilized the *Cell Centered Neighborhoods* function in CytoMAP, which instead of raster scanning neighborhoods, generates neighborhoods centered on the positions of the selected objects. This analysis approach provides a more focused interrogation of the cellular relationships for the selected population of interest. Here, 20 μm radius spherical neighborhoods were centered on the CD31^+^ vascular objects to selectively identify vascular-associated myeloid cell populations (Fig. 7b). This *Cell Centered* approach is demonstrated in Fig. 7b, in which white dots demarcate the centers of all blood vessel objects within the zoom image, and with the example 2D projections of the spherical neighborhoods shown to surround several distinct vascular objects (yellow dots). Once generated, these neighborhoods were clustered as before using the SOM algorithm, with the number of regions determined using the Davies-Bouldin criterion (Fig. S7a). This clustering separated the vascular neighborhoods into several distinct region types (Fig. 7c, top color bar, S7b), which were next manually grouped into four major blood vessel phenotypes based on the local DC subset composition (Fig. 7c, bottom color bar). This clustering revealed that, while many neighborhoods were not closely associated with any given myeloid cell type (unassociated), large portions of the vascular neighborhoods were preferentially associated with either the resident cDC1 or the cDC2 subset. A smaller number of neighborhoods were also associated with CD207^+^ migratory cells (Fig. 7c, S7c). Importantly, 3D spatial remapping of the different vascular subtypes in CytoMAP revealed that large segments of the vascular branches were almost exclusively associated with either the resident cDC1 or cDC2 subset, with little local spatial intermixing (Fig. 7d, Video S2). This indicates that the spatial segregation of LN-resident DC subsets leads to the generation of discrete segments of vascular branches defined by local myeloid cell partners. Thus, in addition to providing potential mechanisms of cDC1 and cDC2 positioning within LNs (based on vascular guidance cues), these data also reveal possible mechanisms for the heterogeneity of LN blood endothelial cells, as recently described.^37,38^

**Figure 7.**
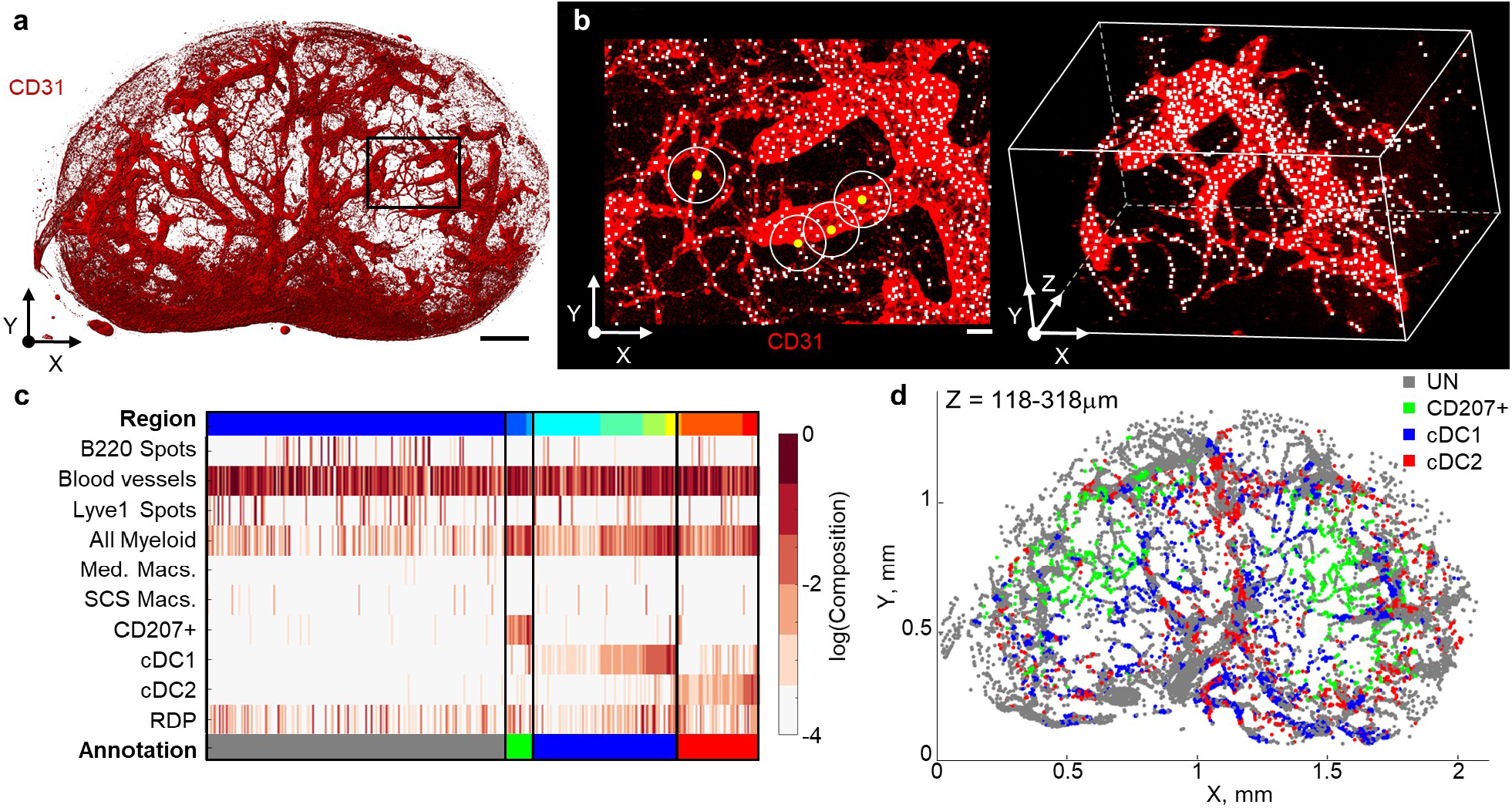
Individual blood vessel branches in LNs demonstrate selective interactions within specific DC subsets. **a**, Confocal image demonstrating the CD31-labeled vascular networks in the same representative LN sample as presented in Fig. 6a. Scale bar = 200μm. **b**, Zoom-in of the region in panel a (denoted with a rectangle), demonstrating the centers of CD31^+^ blood vessel objects (white dots). The circles with yellow dots represent spherical object-centered neighborhoods with a radius of 20μm. Scale bar is 20μm. **c**, Heatmap of the myeloid cell composition for the vessel-centered neighborhoods after SOM clustering. The top color bar shows color-coded regions of neighborhoods. The bottom color bar shows manually annotated regions composed of different DC subsets. **d**, Positional plot of the vascular neighborhoods in a 200μm thick virtual Z section, also color-coded based on the annotations in panel c. Data represent four samples from two independent experiments.

## DISCUSSION

The importance of quantitative imaging and spatial analysis has emerged across a diverse spectrum of biological disciplines at different length scales; from the localization of single molecules within individual cells to the organization of cells across whole organs. Various technologies now allow spatially resolved high-content detection of diverse probe types, from protein and oligonucleotide imaging to mass spectroscopy enabled lipid visualization. These approaches are providing an unprecedented level of detail into biological processes, and as the imaging area and number of analytes increases, the development of tools for analyzing these increasingly complex and voluminous datasets is critical.^4,10,13,14,19,26,30,39,55–57^ Here we developed and described a comprehensive analysis platform, CytoMAP, capable of robust spatial analysis of cellular organization within tissues. CytoMAP harnesses the power of unsupervised clustering, dimensionality reduction, and advanced data visualization to expand the utility of spatially resolved cellular profiling. CytoMAP integrates data on cellular phenotypes and positioning to identify unique neighborhoods and regions within tissues and organs, which provides the ability to interrogate complex spatial patterns across heterogeneous samples. While our technology is demonstrated here using histo-cytometry generated data on cell positioning, CytoMAP in principle can be utilized with datasets generated by diverse approaches and for analysis of various target structures at different length scales. Here we utilized CytoMAP to explore and quantify the organization of immune cells and microenvironments within lymphoid tissues, as well as in tumor and Mtb-infected lung samples. Our results indicate a robust ability of CytoMAP to reveal well-established and previously unappreciated cellular organization within these tissues. In particular, our studies identified the localization of SIRPa^−^ migratory dDCs within the lower cortical ridge of the T cell zone bordering the LN medulla, as well as reveal preferential associations of LN-resident DC populations with blood vessel networks in distinct LN compartments.

In contrast to other approaches, CytoMAP substantially reduces data complexity in two major ways. First, CytoMAP treats complex cell objects as individual points, each possessing information on the positional, morphological and phenotypic cellular characteristics averaged over their respective 3D segmented cellular bodies. Second, CytoMAP defines raster scanned, or cell-centered, local neighborhoods across the tissue, effectively binning data on many similarly positioned cells into single data points. Biologically, such neighborhood binning is equivalent to the concept that, much like urban environments, tissues are subdivided into local neighborhoods containing specific interacting cell populations. Computationally, both steps dramatically reduce the data size (~ 3 orders of magnitude), allowing rapid comparative analysis of cellular composition within the neighborhoods and investigation of how these tissue building blocks are spatially patterned across large 2D and 3D samples. The ability to manipulate the size of the neighborhoods also facilitates exploration of overarching tissue structure or detailed investigation of the hyperlocal cellular microenvironments and cell-cell interactions. Using SOM data clustering, similar neighborhoods are grouped together via topological analysis, which computationally defines unique tissue regions within highly complex datasets with minimal user input. Once clustered, CytoMAP provides tools for exploring the cellular composition and relative prevalence of the tissue regions within and across samples, as well as for visualizing these regions in 2D or 3D space. Interaction network maps provide additional detail into how the regions are spatially interconnected with one another to generate global tissue structure. Additional dimensionality reduction tools allow the user to reveal cellular patterning across the tissues, as well as examine the heterogeneity of individual or multiple samples. Finally, neighborhood based correlation analysis facilitates exploration of how different cell populations are spatially correlated with one another, revealing local cellular interactions, or mutual exclusivity of different cell types with one another. Together, the combination of comprehensive and flexible analytical approaches built into a user-friendly interface makes CytoMAP a powerful toolbox for exploration of complex cellular spatial relationships within large multiplexed imaging datasets.

One area where new spatial analysis tools may provide substantial benefit is in cancer research, in which isolated tumor biopsies have poorly understood cellular organization but still possess substantial prognostic value. To test CytoMAP with such heterogeneous tissues, we explored the organization of immune cells in a CT26 tumor sample. This analysis identified several hallmark features of tumor architecture, including a lymphocytic cuff, as well as centralized positioning of TAMs, which corroborates previous histological studies.^30^ CytoMAP also revealed moderate-to-high baseline infiltration of CT26 tumors by effector T cells, suggesting that this cancer model should be susceptible to checkpoint blockade therapy, which is in line with published data.^47,48^ Interestingly, we found that different myeloid cell types lie in distinct regions within the tumor, suggesting additional layers of cellular organization that should be explored in future studies. Visualizing these relationships across whole tumor cross-sections also revealed substantial intra-tumoral heterogeneity with respect to the local composition of myeloid cells and lymphocytes. This suggests that accurate risk/prognostic assessment of neoplastic tissues may benefit from access to larger tissue samples in addition to punch core biopsies.

As an additional test of CytoMAP, we analyzed cellular organization within granuloma structures in a murine Mtb-infected lung sample. We observed partial segregation of infiltrating CD4^+^ T cells from Mtb-infected myeloid cells, and formation of B cell aggregates. Presence of these distinct tissue regions as identified by CytoMAP’s clustering algorithm largely agrees with previous studies describing immune cell organization in Mtb granulomas, suggesting that CytoMAP presents a promising avenue for investigating the spatial organization of cells in inflamed and infected tissues.^32^

As a final test of CytoMAP, we analyzed the organization of myeloid cells in steady state murine LNs. Consistent with previous observations, we found extensive spatial segregation and a high degree of mutual exclusivity for many of the DC subsets within the LN. In addition, we identified the spatial distribution of a recently described SIRPa^−^ migratory dDC population, which, to our knowledge, has not been positionally mapped previously.^33,34^ We found that these migratory dDCs are predominantly localized within the lower cortical ridge bordering the LN medulla, and together with the locally positioned SIRPa^+^ dDCs generate a prominent cuff-like cellular aggregate. In addition, analysis of resident cDC1 and cDC2 organization recapitulated previous findings, with preferential localization of resident cDC2 in the LN periphery and more heterogeneous distribution of resident cDC1s across the T cell zone and reduced presence in the LN medulla.^4,5,7–9,52,58^ Importantly, we found that both resident cDC1 and cDC2 subsets were highly associated with LN blood vessels, albeit preferentially associating with distinct vascular trees, with little local intermixing. Although previous studies have shown that DCs can associate with blood vessels during inflammation,^59,60^ our findings reveal that this normally occurs in the steady state, and may thereby promote homeostatic maintenance of LN blood vessels.^35,36^ Our studies also suggest that blood vessels could provide positional cues to guide resident cDC1 and cDC2 distribution in LNs. While previous studies have identified a role for the G-protein coupled receptor 183 in guiding resident cDC2 positioning and survival in lymphoid tissues,^61,62^ little is known about the regulation of resident cDC1 localization in LNs. Our study provides hints to a potential mechanism regulating cDC1 distribution, and will require further study. Finally, our studies suggest that DC subset spatial patterning and exclusive interactions with distinct vascular branches may also influence blood endothelial cell biology, potentially promoting the recently described heterogeneity of LN blood endothelial cells.^37^

In sum, here we develop a user-friendly, comprehensive, and broadly applicable spatial analysis toolbox for analysis of 2D or 3D quantitative imaging datasets, which excels at utilizing high-dimensional imaging data to reveal complex tissue features based on cellular phenotypic heterogeneity and spatial patterning. Our technology allows robust cross-sample interrogation of spatial cellular relationships, tissue architecture, and reveals intra- as well as inter-sample heterogeneity. In this early implementation, CytoMAP has already provided new insights into the organization of myeloid cells in lymphoid tissues, revealing the localization of SIRPa^−^ migratory dDC, as well as identifying preferential associations of resident DCs with select LN vasculature. Phenotyping neighborhoods with unsupervised clustering revealed distinct regions which were both biologically relevant and consistent across multiple samples. Together, this indicates that CytoMAP is a valuable tool for the rapid identification of key cellular networks and tissue structure, revealing the fundamental building blocks of tissue organization.

## ACKNOWLEDGMENTS

We thank Dr. Frank Schmitz (Celgene/Fred Hutch) for many useful discussions. We additionally thank the open source software community, including HistoCat, Flow Cytometry GUI for Matlab, fca_readfcs (fca 2.2). This work was in part funded by Roche Glycart, the Washington Research Foundation postdoctoral fellowship (CS), and NIH grants R01AI134713 (MYG), R21AI142667 (MYG), 1R01AI134246 (KBU), 1R01AI076327 (KBU), U19AI135976 (KBU), T32HD007233 (BHG).

## AUTHOR CONTRIBUTIONS

CS and MG analyzed data and prepared manuscript. CS and JF wrote the software. CS and MG conceived of the project and software. CS, BO, JL, BG, MG, and CP performed mouse studies and imaged samples BO and MG Validated Antibodies. MP and TP provided tumor samples. BG, CP, and KU provided lung samples. CS, JF, BO, BG, MLC, BEO, JL, CPS, TP, MP, JH, and MG provided software feature, design, and usability feedback. CS, CPS, MG, and JF conceived of and implemented statistical approaches. MG supervised the project. All authors edited manuscript.

## DATA AVAILABILITY

CytoMAP software is available for download at https://gitlab.com/gernerlab/cytomap. Imaris extensions and other code used for histo-cytometry analysis is available for download at: https://gitlab.com/gernerlab/imarisxt_histocytometry. All data are available upon request.

## Materials and Methods

### Mice

For the experiments described in Fig. 2, 5-7, 6-10 week old male and female B6.SJL and C57BL/6J mice were obtained from The Jackson Laboratory and kept in specific pathogen–free conditions at an Association for Assessment and Accreditation of Laboratory Animal Care–accredited animal facility at the University of Washington, South Lake Union campus. All procedures were approved by the University of Washington Institutional Animal Care and Use Committee.

For the data presented in Fig. 3, Balb/c mice were obtained from Charles River (Sulzfeld, Germany) and were housed in specific pathogen-free conditions. The animal facility was accredited by the Association for Assessment and Accreditation of Laboratory Animal Care (AAALAC) and all animal studies were performed in accordance with the guidelines outlined by the Federation for Laboratory Animal Science Association (FELASA) and the German Animal Welfare law. The animal study was approved by and done under license obtained from the Government of Upper Bavaria (Regierung von Oberbayern; license number: ROB-55.2-2532.Vet_03-15-41).

For the experiment described in Fig. 4, C57BL/6J mice were obtained from The Jackson Laboratory and housed in specific pathogen-free conditions at Seattle Children’s Research Institute (SCRI). Experiments were performed in compliance with the SCRI Animal Care and Use Committee. An 8 week old female mouse was used for the presented study.

### LN studies

For the experiment shown in Fig. 2, a C57BL/6 mouse was adoptively transferred with 10^6 naïve OT-II CD4+ T cells and one day later immunized in the footpad with OVA plus Alum; 4.5 days later the draining popliteal LN was harvested and used for analysis. For the experiments shown in Fig. 5, non-draining steady state skin LNs were harvested from C57BL/6 mice which were previously injected in the contralateral distal footpad with Alum 2 days before harvest. For the experiments shown in Fig. 6 and 7, skin LNs were harvested from naïve C57BL/6 mice.

### Tumor studies

Balb/C mice were injected subcutaneously with 5×10^6 CT26.WT cells and 9 days later, the tumor was harvested for fixation and imaging.

### Mtb studies

All infections were done with a stock of Mtb H37Rv, as previously described.63 To perform aerosol infections, C57BL/6 mice were enclosed in a Glas-Col aerosol infection chamber, and Mtb bacilli were deposited directly into their lungs. Lungs were removed 34 days post infection.

### Tissue preparation – thin sections

All thin tissue sections were fixed with Cytofix (BD Biosciences) buffer diluted 1:3 with PBS for 12h at 4° C and then dehydrated with 30% sucrose in PBS for 12-24h at 4° C. Tissues were next embedded in O.C.T. compound (Tissue-Tek) and stored at −80° C. LNs were sectioned on a Thermo Scientific Microm HM550 cryostat into 20μm sections and were then prepared and imaged as previously described.^4^ Briefly, sections were stained with panels of fluorescently conjugated antibodies, shown in table S5, cover-slipped with Fluoromount G mounting media (SouthernBiotech), and imaged on a Leica SP8 microscope.

### Tissue preparation – thick slices

Volumetric imaging using Ce3D tissue clearing (thick sections) was performed as previously described.13,39 In brief, LNs were fixed with Cytofix (BD Biosciences) buffer diluted 1:3 with PBS for 12-20h at 4° C. Excess fat was carefully removed using a dissection microscope, and the samples were embedded in 2% Agarose. 500um thick cross-sectional slices were generated using a Vibratome (Leica VT1000S, Speed: 215 Frequency: 8). Slices were next placed in blocking buffer (1%NMS, 1%BSA, 0.3%Triton, in 0.1MTris) for 24h at 24° C on a rocker. After blocking, LN slices were stained with a panel of directly conjugated antibodies (table S5) for 3 days at 34° C on a shaker, then washed in blocking buffer for one day at 24° C. Next, slices were placed in Ce3D clearing solution (13.75ml 40% [vol/vol diluted with PBS] N-methylacetylamide; 20g Histodenz; 25uL Triton X-100; 125ul Thioglycerol) at 24° C for at least 24h. Finally, slices were cover-slipped using Ce3D as the mounting media and imaged on a Leica SP8 microscope.

### Imaging

All samples were imaged using a Leica confocal SP8 microscope, with either a 40X 1.3NA (HC PL APO 40×/1.3 Oil CS2, for 20μm sections) or a 20X 0.75NA (HC PL APO 20×/0.75 IMM CORR CS2, free working distance = 0.68 mm, for thick cleared slices) oil objective with type F immersion liquid (Leica, refractive index n_e_ = 1.5180). After acquisition, stitched images were compensated for spectral overlap between channels using the Leica Channel Dye Separation module in the Leica LASX software. For single stained controls, UltraComp beads (Affymetrix) were incubated with fluorescently conjugated antibodies, mounted on slides, and imaged with the same microscope settings used in the image they were being used to compensate.

### Image analysis and histo-cytometry

Image analysis and Histo-cytometry was performed as described previously, with minor modifications.^4,13,39^ A detailed description is available in the supplemental information.

### Statistical analysis

No statistical method was used to predetermine sample size. The statistical significance of Pearson’s correlation was calculated using a Student’s t distribution for a transformation of the correlation.

### CytoMAP spatial analysis

CytoMAP was written using MATLAB version 2018b (Mathworks). A detailed description of the workflow and functions built into CytoMAP is available in the online user manual. A brief discussion of the analyses used in this manuscript is described in the supplemental information.

## DATA AND CODE AVAILABILITY

Imaris extensions and other scripts used for histo-cytometry analysis are available for download at: https://gitlab.com/gernerlab/imarisxt_histocytometry CytoMAP software is available for download at: https://gitlab.com/gernerlab/cytomap

