## Supplementary Information for "CytoMAP: a spatial analysis toolbox reveals features of myeloid cell organization in lymphoid tissues"

***Image analysis and histo-cytometry***

Histo-cytometry was performed as described previously, with minor modifications. Briefly, using Imaris, various steps were taken to process the images and segment the cell objects. Unless otherwise noted, only the channel arithmetic and surface creation steps for each sample shown in tables S2 and S3 were used to process the images. Channel arithmetic were performed using either the default Imaris function or a customized ImarisXT extension, *Calebs_Multi_EQ_ChannelArithmetics_V3* (available online), which allows for batching multiple equations and auto saving the image once the function is done running. Additional normalization and processing steps for specific samples are described below. After surface creation, the MFI for each imaged channel, as well as the volume, sphericity, and position of the cell objects were exported and concatenated into a single .csv file using the *Imaris_To_FlowJo_CSV_Converter_V4* MATLAB function, which is a customized .csv merge tool available online. The combined .csv file was next imported into FlowJo (TreeStar) and the cell objects were classified into the indicated cell subsets according to the gating strategies shown in the respective figures.

For the experiments described in Fig. 2, using Imaris version 8.3 the All Cell Composite channel was created using the equation in table S2. Using the surface creation tool in Imaris, masking surfaces were created around this new channel using the parameters for surface_1 shown in table S3. Using the mask option, all pixels from the new channel outside the boundaries of this surface were set to equal 255 (the maximum pixel intensity). Next the new channel was smoothed with a Gaussian filter with a width of 0.56µm. This new channel was then inverted, and the gamma was set to 0.3. Finally, cell object surfaces were created using the inverted new channel with the properties shown in table S3 for surface_2.

For both experiments shown in Fig. 6, all image layers were first normalized along the Z axis. For experiment 1, a clean CD31 channel was created to correct for residual channel spillover from the Lyve1 channel. Next a composite channel of myeloid cell markers was created according to the equation in table S2. Finally, surfaces were created according to the parameters in table S3. For the images from the second experiment, channels SIRPa, B220, MHC-II, CD31, and CD11c were smoothed using the Imaris *Gaussian Smooth* function with a width of 0.56µm. Next, a composite channel of myeloid cell markers was created according to the equation in table S2. Finally, surfaces were created according to the parameters in table S3.

For the image of the lung shown in Fig. 4, a composite of all surface markers was created using the equation in table S2. The linear stretch function in Imaris was used to set the blackpoint of this channel to 40, gamma was adjusted to 0.75, and Gaussian smoothing was used with a width of 0.32um. The Jojo1 nuclear stain channel was brightened with the linear stretch function in Imaris to set the saturation point to 200, gamma was adjusted to 1.5, and Gaussian smoothing was used with a width of 0.32um. The all surface marker channel was then subtracted from the altered Jojo1 channel to form a "Nuclei" channel according to table S2. Finally, surface objects were created in Imaris on this new channel using the parameters outlined in table S3.

***CytoMAP spatial analysis***

Below is a brief discussion of the analysis used for the datasets described in this manuscript. The annotated cell surfaces for each dataset were loaded into CytoMAP by importing the corresponding paired .wsp and .fcs files after saving them in FlowJo. These collectively contain the cell statistics, gate definitions, and spatial positions for each cell object. Once imported the following functions were used in CytoMAP to analyze the data.

#### Generate Random Points

This function, found in the ribbon at the top of the figure window, uses MATLAB's built in *rand* function, with all default parameters, to generate a set of random points based on the set of currently plotted points in the *New Figure* window. This function only generates randomized data with respect to the axes which are plotted, while setting all other channel values to 0 for the new set of points. Additionally, this function scales the randomized values to be within the limits of the currently plotted points. To generate points with a random spatial distribution, all cells were first plotted with positions on the X and Y axes. Next, the Generate Random Points function was called and this newly generated set of points was saved. Finally, all points outside the spatial bounds of the tissue were discarded by gating on objects within the tissue boundaries, resulting in sets of RDPs for each sample analyzed, with a representative set of points shown in Fig. S4b.

#### Make Surface

This function uses MATLAB's *alphaShape* function, with user defined parameters, to wrap the currently plotted points in a surface. This surface can then be used to gate on points, such as randomly distributed points, and to calculate the distance to region borders. This function was used to generate the surfaces shown in Fig. 5f.

#### Calculate Distance

This function finds the physical distance, either to the border of user defined surfaces, or to the nearest cell. This function was used to find the distance to region borders plotted in Fig. 5 g-j, and the distance between myeloid cells and the nearest blood vessel plotted in Fig. 6b.

#### Raster Scan Neighborhoods

This function uses Neighborhood analysis, which finds the local composition of cells within a circular (2D data) or spherical (3D data) area/volume in the tissue. If the data has non-zero z dimensionality but the z thickness is less than the radius of the neighborhoods, this function will treat the data as effectively 2D and use a cylindrical neighborhood window. This function calculates the number of cells and the MFI of each channel summed over all cells in each neighborhood. The positions of the neighborhoods are evenly distributed throughout the tissue in a grid pattern with a distance between neighborhood centers of half of the user defined radius. The neighborhood information can then be used for further analysis (e.g. local cellular densities, cell-cell associations).

#### Cell Centered Neighborhoods

This function is similar to *Raster Scan Neighborhoods*, except the position of the neighborhoods are centered on a user selected cell type. Once the cell type is chosen and the radius is defined, this function calculates the number of cells and the MFI of each channel summed over all cells in each neighborhood.

#### Classify Neighborhoods Into Regions

This function is used to define tissue regions using multiple types of information about the neighborhoods. This includes, the standardized number of cells of each phenotype in each neighborhood (number of cells minus mean number of cells, divided by the standard deviation of the number of cells in each neighborhood across the dataset), composition (number of cells divided by the total number of cells in each neighborhood) and raw number of cells per neighborhood, which were used as indicated in the respective figures. In this manuscript, the physical position of the neighborhoods was not used for region definition and the minimum of the Davies-Bouldin function was used to automatically determine the number of regions. The neighborhoods were clustered using the SOM function. This utilizes MATLAB's *selforgmap* function with default parameters, except the dimensions options, which is equal to the number of regions (NR) by one, i.e.

app.net.(ModelName).Network = *selforgmap([NR, 1])*

Once the network parameters are defined, the network is trained on the neighborhood data using MATLAB's *train* function. This assigns a cluster number to each neighborhood. The *selforgmap* algorithm starts with NR "neurons" positioned throughout the data. It then iteratively moves the position of the neurons closer to the data to match the landscape of the data. Here, the position is not the spatial position, but the topological position within the cell composition data. The neighborhoods are then clustered by finding the closest neuron for each neighborhood. For visualization purposes the arbitrarily color designations of the individual regions were changed using the *Edit Region Colors* function in CytoMAP. The composition of the color-coded neighborhoods was plotted using the *Heatmap Visualization* function in CytoMAP. The spatial distribution of the regions was visualized by generating a new figure in CytoMAP, plotting the positions of the neighborhoods, and selecting the regions for the 'c' axis to color-code the neighborhoods by region type.

#### Reduce Dimensions

This function uses the built in MATLAB implementation of the t-SNE algorithm to reduce the neighborhood data to two dimensions. For the t-SNE plots in this manuscript the default MATLAB options were used, including: Euclidian distance, perplexity of 30, theta, of 0.5, and exaggeration of 4.

#### Pseudo-Space

This function allows the user to sort the neighborhoods by the absolute number or composition of different cell types within the neighborhoods and plots the neighborhoods in this sorted order on a linear pseudo-space axis. The Y axis is normalized to allow comparison between cell types, and the data are smoothed along the X axis by a user defined amount. This function was used with the data type, weights, and smoothing parameters for each figure shown in table S4.

#### Cell-Cell correlation analysis

This function calculates the Pearson correlation coefficient of the number of cell or object types within the scanned neighborhoods and graphs these on a heatmap plot.

#### Network Map

This function calculates the percentage of neighborhoods of each region type which are directly in contact with each other region type. It creates a force directed graph with a node for each region type, where the nodes are connected by an edge if more than 0.005% of the neighborhoods of that nodes type are in contact with the connecting node's region type. The edge thickness is proportional to the % of neighborhoods in contact with the connecting node region type, and the node size is proportional to the number of neighborhoods of the region type.

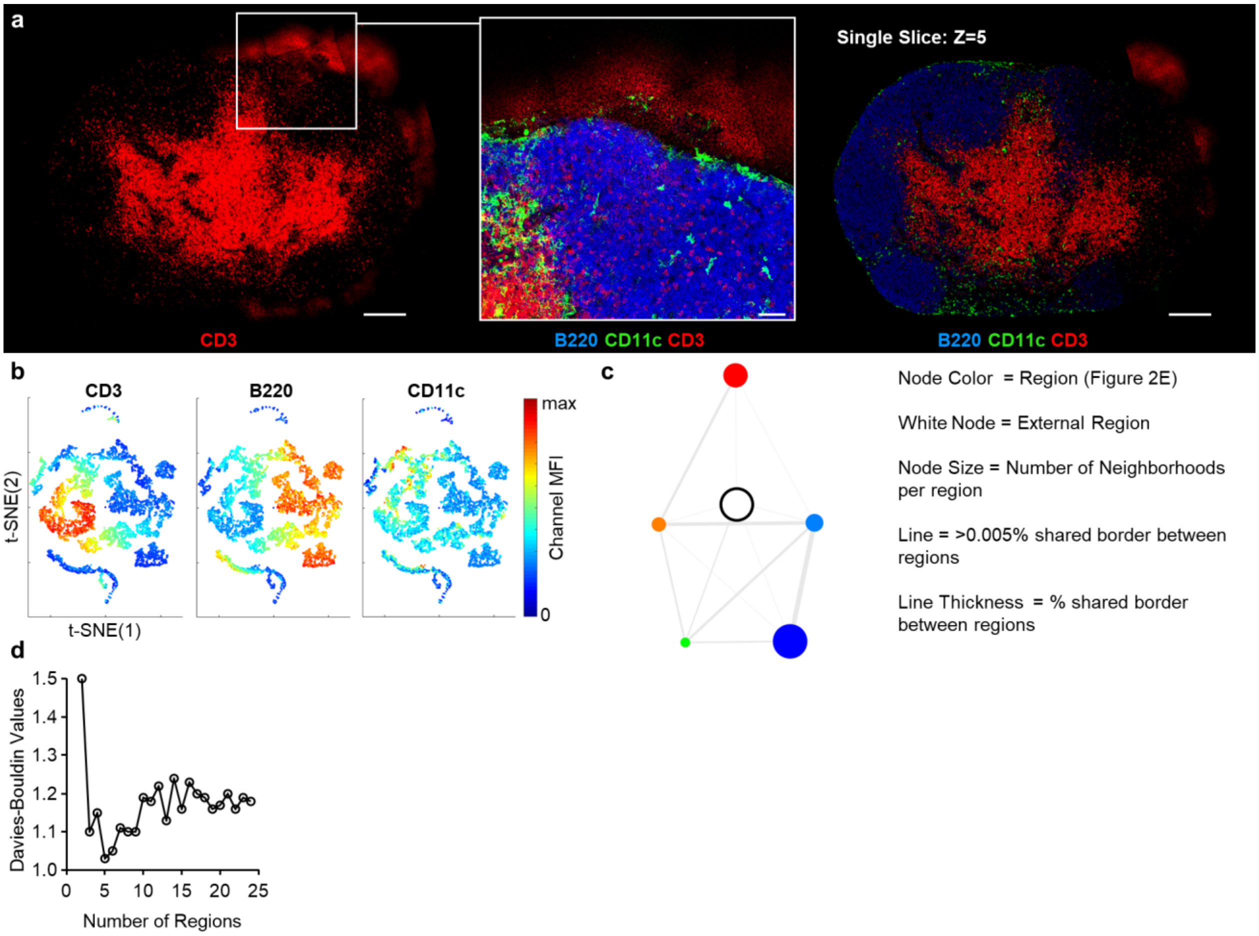


**Figure S1. Identification of basic LN architecture with CytoMAP - related to Fig. 2. a**, The same LN dataset as presented in Fig. 2a, demonstrating the out-of-plane noise in the CD3 channel within the maximum projection image (left) or a single Z-slice (right). Overview image scale bar = 200µm; zoom-in scale bar = 50µm. **b**, Extended t-SNE analyses for Fig. 2g, demonstrating a color-coded heatmap of the total MFI signal in each neighborhood for the indicated markers. **c**, Interaction map showing the percentage of shared border between the regions, as defined and color-coded in Fig. 2e. The white node represents area external to the tissue. The node size is proportional to the prevalence of that region within the sample (number of neighborhoods per image). Lines connect regions that share borders, with the line thickness being proportional to the percent of the border shared between the regions. **d**, Davies-Bouldin values were used to determine the number of regions presented in Fig. 2d.

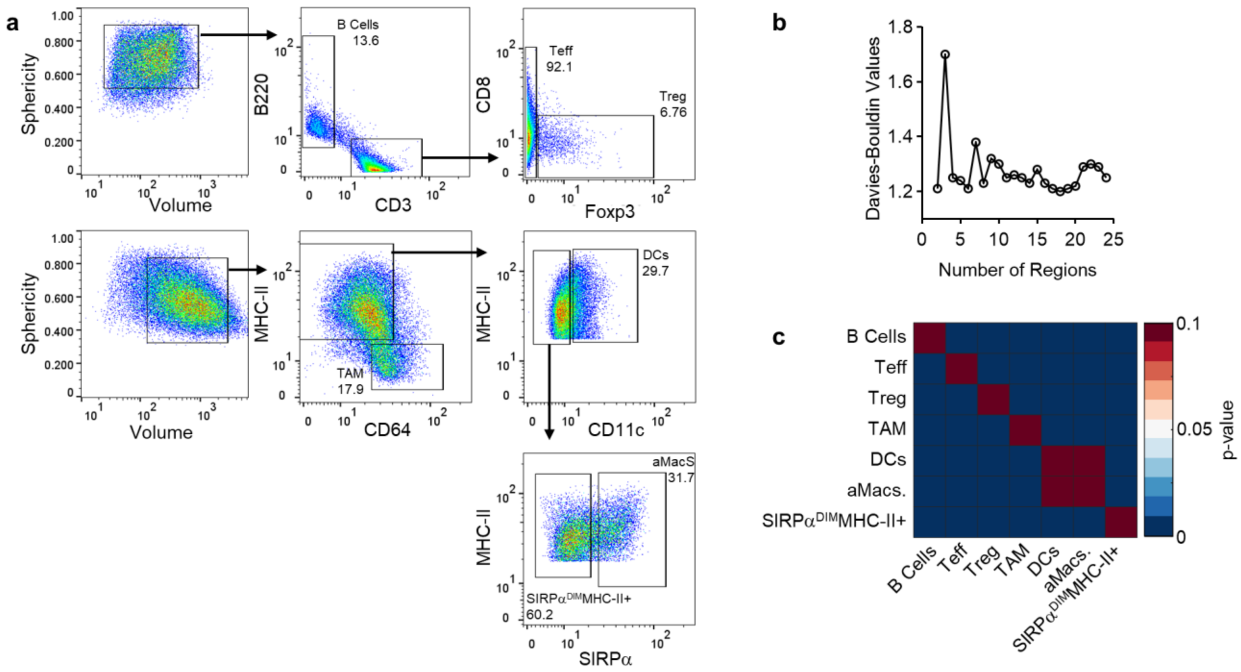


**Figure S2. Quantitative analysis of tumor immune infiltrate – related to Fig. 3. a**, Histo-cytometry plots of the gating strategy used to annotate the cell types for the sample analyzed and presented in Fig. 3. **b**, Davies-Bouldin values used to determine the number of regions in the neighborhood dataset, with the SOM clustering presented in Fig. 3c. Although the absolute minimum of the function indicated presence of up to 18 different regions, 6 regions were used for clustering, as this was relatively similar to the function minimum. **c**, Heatmap of the p-values for the corresponding Pearson correlation coefficients presented in Fig. 3f. High p-values correspond to correlation coefficients not significantly different from 0.

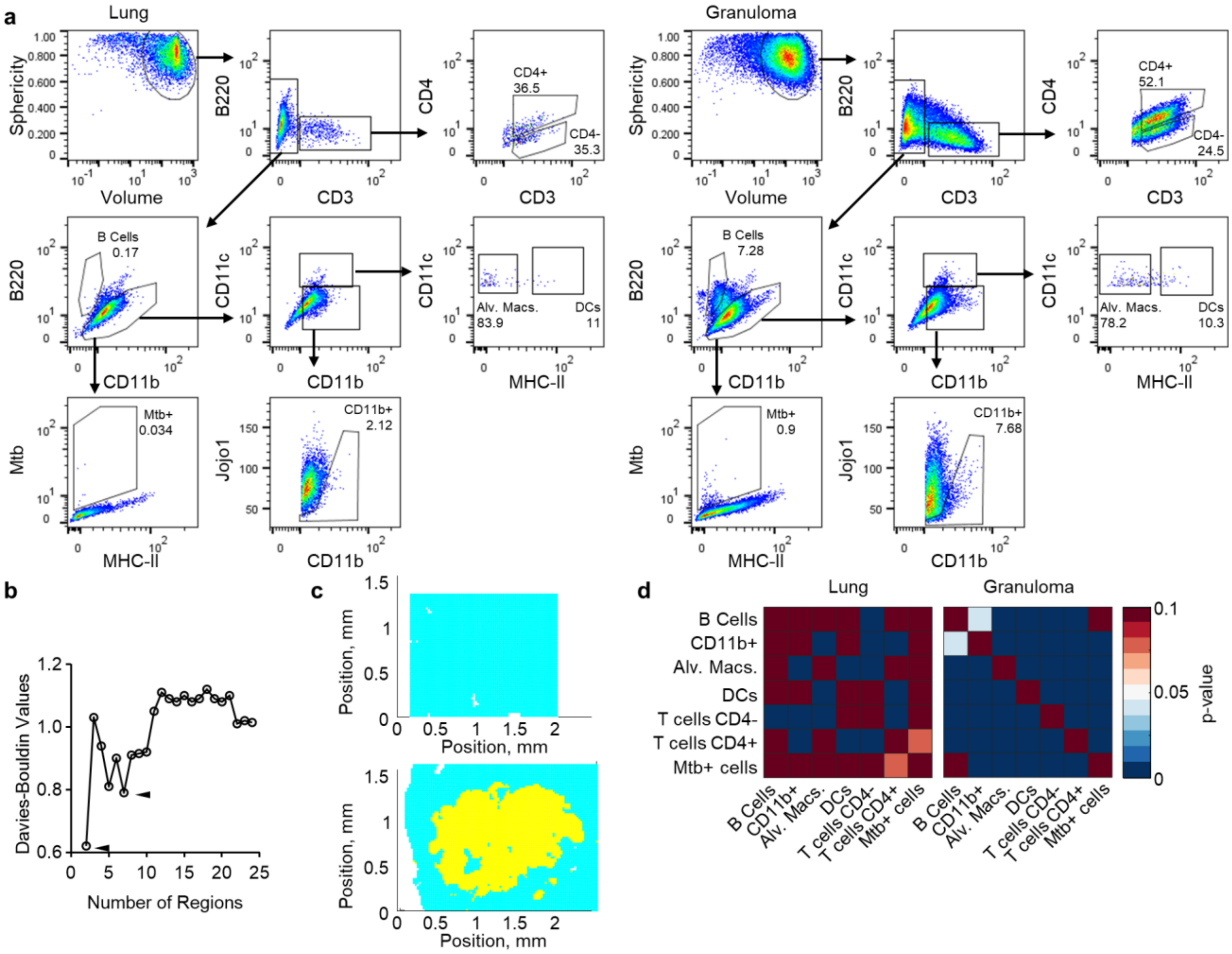


**Figure S3. Quantitative analysis of lung Mtb granulomas – related to Fig. 4. a**, Histo-cytometry plots showing the gating strategy used to annotate the cell types presented in Fig. 4b. The cell objects extracted from the uninvolved lung image are on the left and the granuloma image are on the right. **b**, The Davies-Bouldin values used to determine the number of regions in the neighborhood dataset, with the SOM clustering presented Fig. 4c. The neighborhoods were clustered into 6 regions based on these values. Since there are two very different regions in this dataset (lung and granuloma) the minimum of the Davies-Bouldin yields only two regions. However, the local minimum at 6 regions reveals both the difference between the lung and granuloma neighborhoods, as well as the different regions within the granuloma. **c**, The color-coded position plot of the regions if 2 regions were used based on the absolute minimum from the Davies-Bouldin function, as presented in panel c. This plot demonstrates two distinct regions, the uninvolved lung (cyan) and the granuloma (yellow). **d**, The p-values for the corresponding Pearson correlation coefficients presented in Fig. 4f. High p-values correspond to correlation coefficients not significantly different from 0.

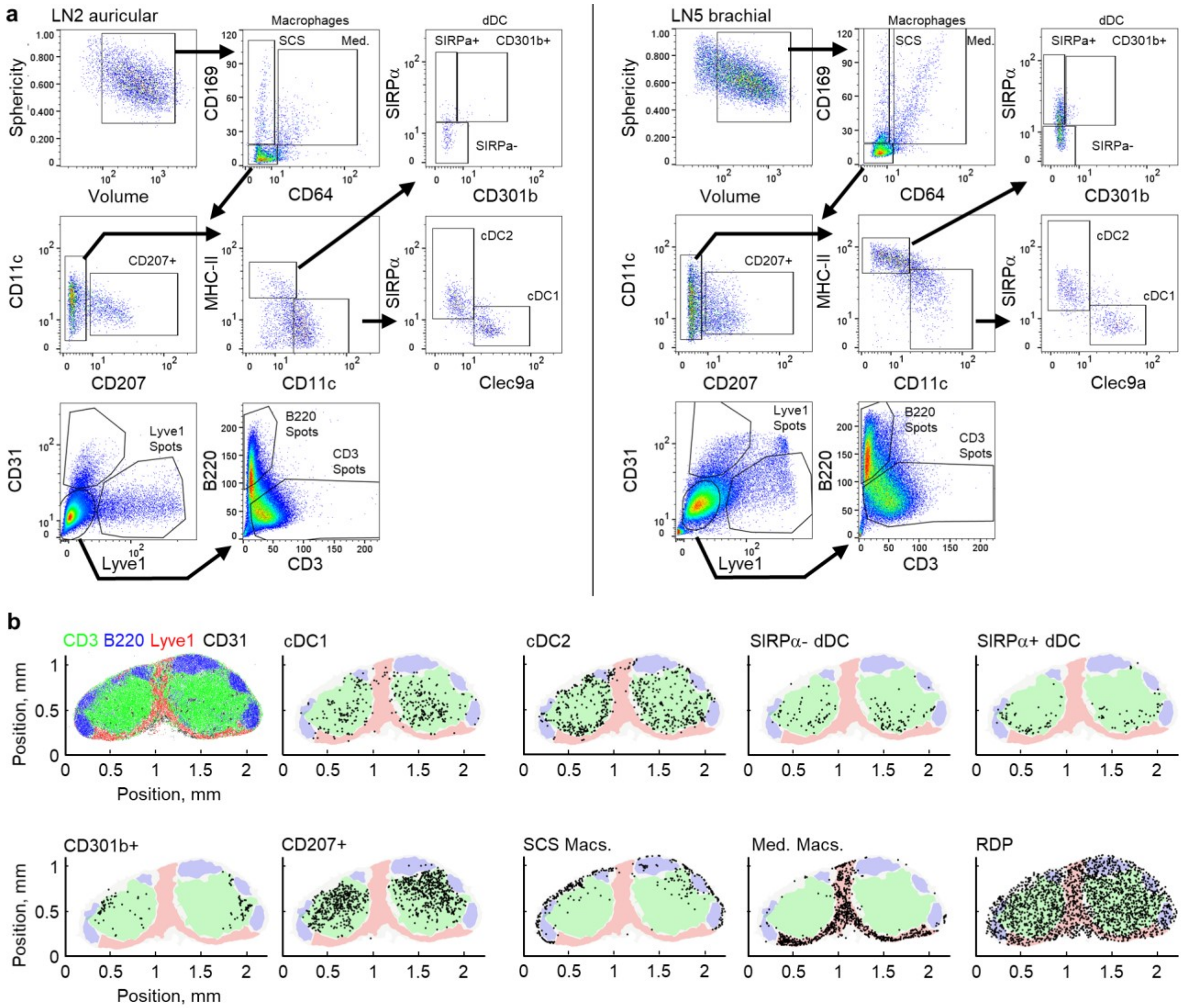


**Figure S4. Quantifying myeloid cell organization in LNs - related to Fig. 5. a**, Two independent representative histo-cytometry plots demonstrating the gating strategy used to annotate cell types and landmark spots in the dataset presented in Fig. 5. **b**, Positional plots of the landmark spots (top left), indicated cell populations, and RDP (bottom right) overlaid on the manually annotated regions, as shown in Fig. 5f.

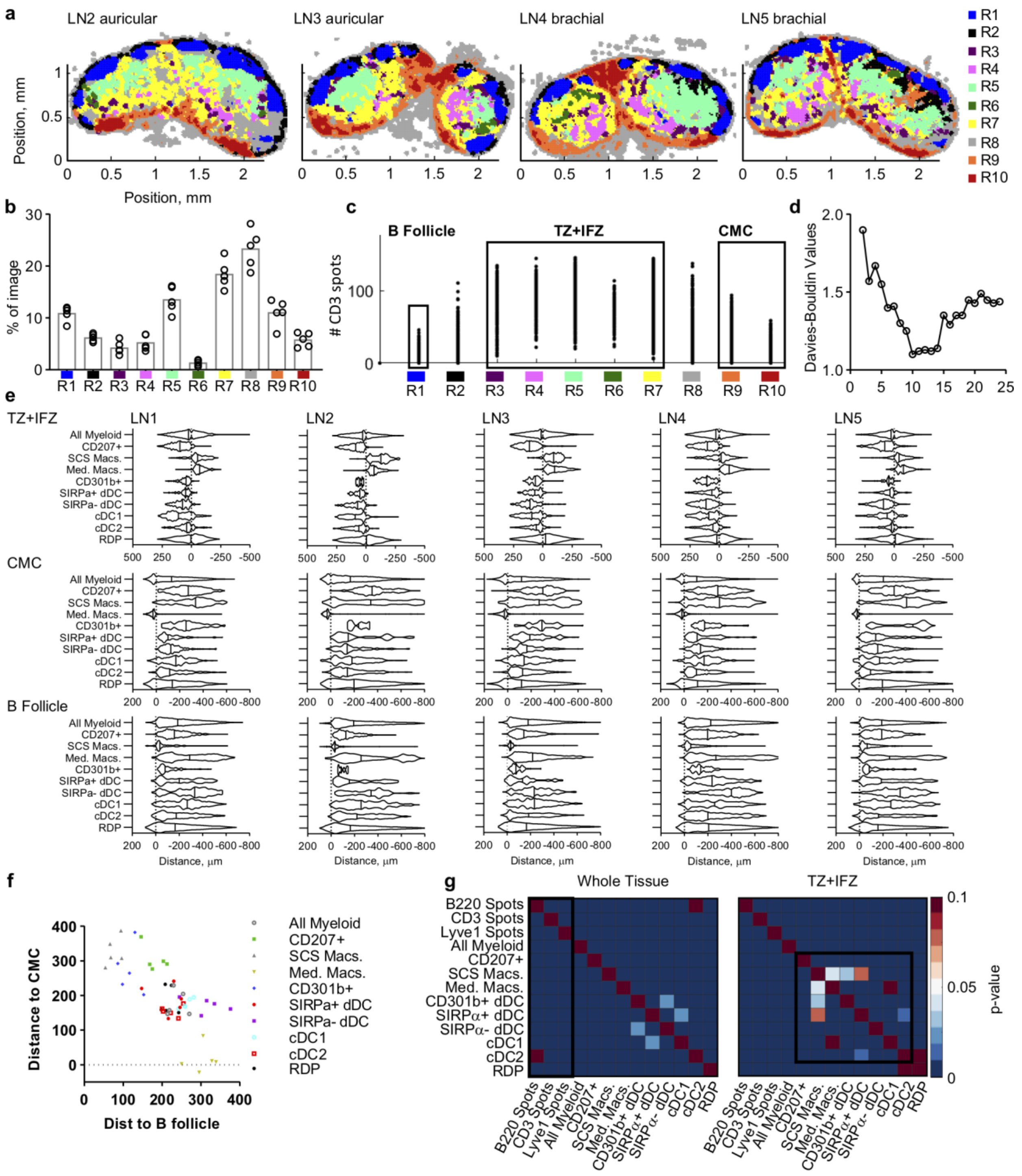


**Figure S5. CytoMAP reveals patterns of myeloid cell organization in LNs - related to Fig. 5. a**, Color-coded spatial plots of the neighborhoods, demonstrating the distribution of CytoMAP generated regions across all of the LNs presented in Fig. 5. **b**, Region prevalence plot showing the percentage of the neighborhoods from each sample in each region. **c**, Representative gates on the neighborhoods based on their region number used to manually annotate the regions and define the surfaces shown in Fig. 5f. **d**, Davies-Bouldin values used to determine the number of regions presented in Fig. 5d. **e**, Violin plots of the distances for all cells from each LN. The means of these distances are shown in Fig. 5h-j. TZ+IFZ distance analysis for LN1 are the same data as presented in Fig. 5g. **f**, Combined analysis for Fig. 5i and 5j. Plot demonstrates the average distance of the indicated cell populations to the B cell follicle versus the CMC, with each dot representing an individual sample. **g**, Heatmaps of the p-values for the Pearson correlation coefficients shown in Fig. 5k. High p-values correspond to correlation coefficients not significantly different from 0. Data represent 5 samples from one independent experiment.

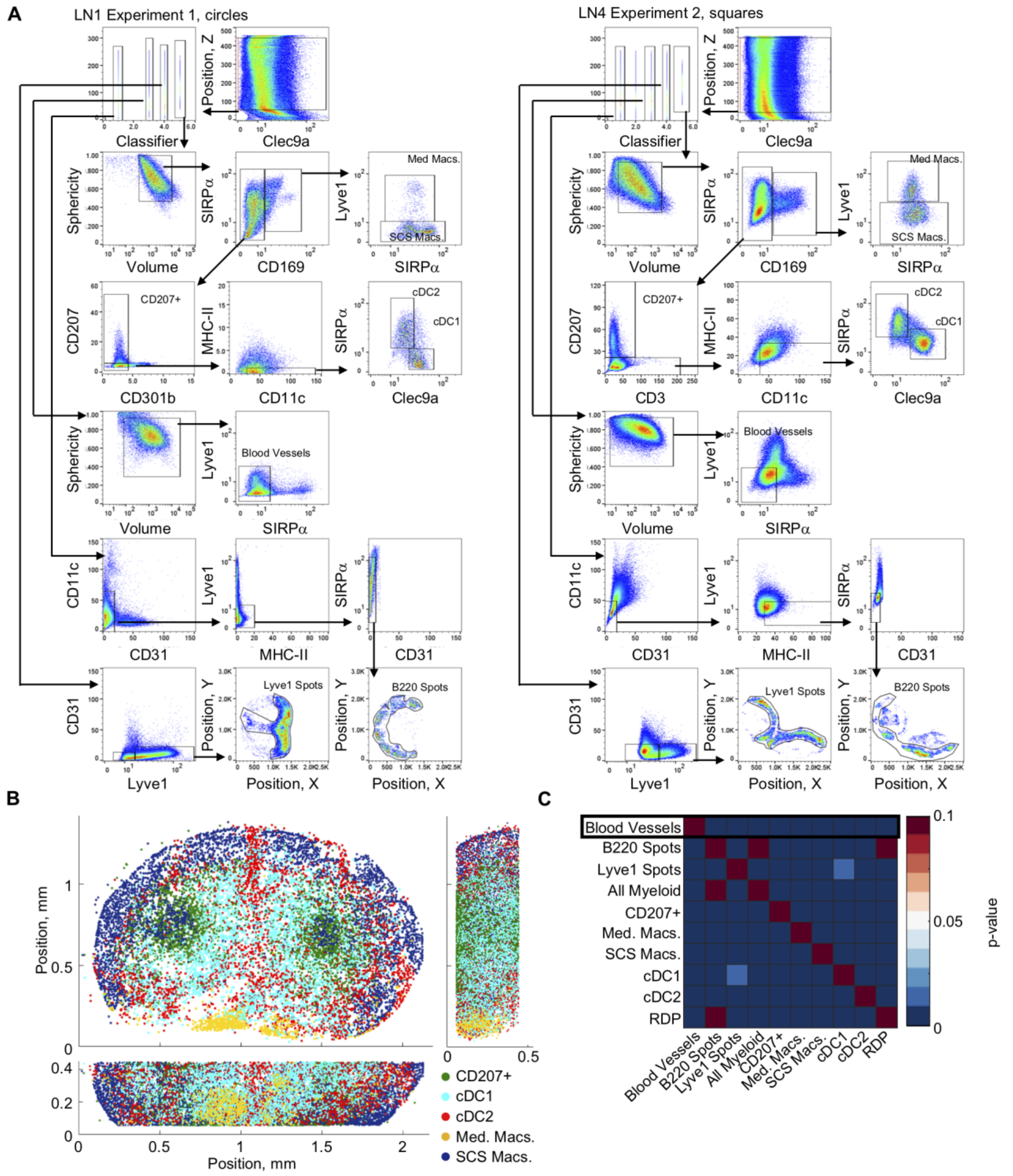


**Figure S6. Identification of DC subsets in 3D LNs – related to Fig. 6. a**, Histo-cytometry hierarchical gating of the cell populations extracted from two representative samples, one for each experiment, for the data presented in Fig. 6. The classifier is a manually added parameter used to distinguish between different sets of surface objects for gating purposes. **b**, 3D spatial remapping of the cell objects annotated in panel a. **c**, Heatmap of the p-values for the Pearson correlation coefficients presented in Fig. 6e. High p-values correspond to correlation coefficients not significantly different from 0. Data represent four samples from two independent experiments.

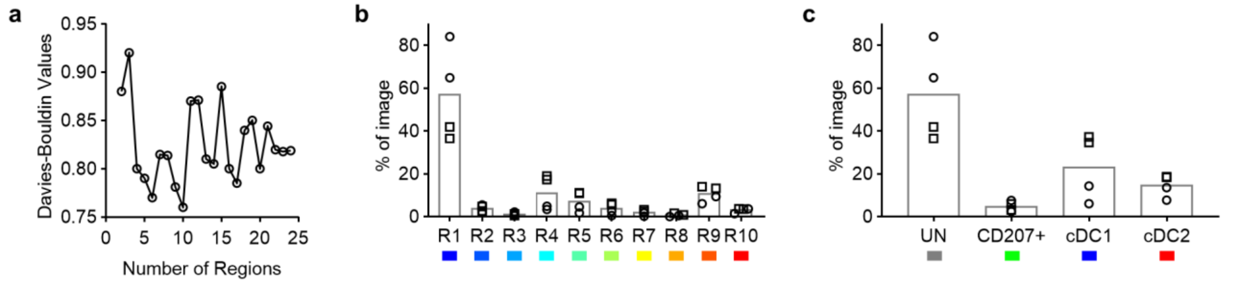


**Figure S7. Characterization of blood vessel branches in LNs – related to Fig. 7. a**, Plot of the Davies-Bouldin values used to determine the number of regions used for SOM clustering presented in Fig. 7c. **b**, Percentage of neighborhoods in each cluster defined in the color bar at the top of Fig. 7c. **c**, Percentage of neighborhoods in each manually defined cluster group from the bottom color bar in Fig. 7c. Data represent four samples from two (circles vs. squares) independent experiments.

**Video S1. Association of resident cDC1 with LN vasculature – related to Fig. 6.** This video shows a 3D visualization of the Ce3D cleared steady state LN (also shown in Fig. 6a), demonstrating close association of Clec9a+ cDC1 with CD31+ LN blood vessels. Clec9a staining is cyan, CD31 staining is red and Lyve1 staining is purple. Clec9a expressing cells are predominantly cDC1 cells.

**Video S2. DC subsets associate with distinct LN vascular trees – related to Fig. 7.** This video shows a 3D view of the vascular networks (also shown in Fig. 7d), demonstrating large segments of vascular trees preferentially associated with distinct DC populations. Each dot represents a vascular neighborhood. Unassociated vascular neighborhoods – grey; CD207 associated neighborhoods – green; resident cDC1 associated neighborhoods – blue; resident cDC2 associated neighborhoods – red.

Table S1. Reagents and Resources

| REAGENT or RESOURCE | SOURCE | IDENTIFIER |
|---|---|---|
| **Antibody-fluorophore (clone)** | | |
| CD3 -BV421; -AF700; -APC-F750; -CF633 [Conjugated in house] (17A2) | BioLegend | Cat# 100228; 100216; 100247; 100202 |
| CD4-CF660 (RM4-5) [Conjugated in house] | BioLegend | Cat# 100506 |
| CD8 (53-6.7)-BV510 | BioLegend | Cat# 100752 |
| CD11b (M1/70) | BD | Cat# 55308 |
| CD11c -BV421; -AF647 (N418) | BioLegend | Cat# 117330; 117312 |
| CD11c-eFluor615 (N418) | eBioscience | Cat# 42-0114-82 |
| CD11c-BV480 (HL3) | BD | Cat# 565627 |
| CD31 -BV421; -BV480 (390) | BioLegend; BD | Cat# 102424; 746260 |
| CD31-AF488 (Mec 13.3) | BioLegend | Cat# 102513 |
| CD45.2-BV510 (104) | BioLegend | Cat# 109838 |
| CD64 -PE; -CF660 [Conjugated in house] (X54-5/7.1) | BioLegend | Cat# 139304; 139302 |
| CD169-CF514 (3D6.112) [Conjugated in house] | BioLegend | Cat# 142402 |
| CD207 -AF488; -AF546 (929F3.01) | Fisher Scientific | Cat# DDX0362A488; DDX0362A546 |
| CD301b-CF660 (URA1) [Conjugated in house] | BioLegend | Cat# 146802 |
| B220 -biotin; -BV510; -AF700; -CF750 [Conjugated in house] (RA3-6B2) | BioLegend | Cat# 103204; 103247; 103232; 103202 |
| B220 (RA3-6B2)-DY405LS | Novus Biologicals | Discontinued |
| CXCR3-PE (CXCR3-173) | Fisher Scientific | Cat# 12-1831-82 |
| Foxp3-eF570 (FJK-16s) | eBioscience | Cat# 41-5773-82 |
| IFNg-PE (XMG1.2) | BioLegend | Cat# 505807 |
| iNOS (C-11)-AF405 | Santa Cruz | Cat# sc-7271 |
| IRF4 -CF514; -CF594 (IRF4.3E4) [Conjugated in house] | BioLegend | Cat# 646402 |
| Jojo1 | Invitrogen | Discontinued |
| Ki67-AF700 (B56) | BD | Cat# 561277 |
| Lyve1 -biotin; -eFluor570 (ALY7) | eBioscience | Cat# 13-0443-82; 41-0443-82 |
| MHC-II -AF700; -Dy395xl (M5/114.15.2) | BioLegend | Cat# 107622; 107602 |
| Mtb (rabbit polyclonal) | Abcam | Cat# Ab905 |
| PCREB-AF488 (87G3) | Cell Signaling | Cat# 9187S |
| pS6 (2F9)-AF488 | Cell Signaling | Cat# 48554 |
| SIRP$\alpha$ -BV421; -CF594 [Conjugated in house] (P84) | BD; Biolegend | Cat# 624124; 14401 |
| TCF1-PacBlue (C63D9) | Cell Signaling | Cat# 9066S |
| anti-rabbit-AF750 | Invitrogen | Cat# A-21039 |

| | | |
|---|---|---|
| SA-AF750 | Invitrogen | Cat# S21384 |
| SA-ATTO490LS | ATTO-TEC | Cat# AD490LS-61 |
| anti-GFP-AF488 | Invitrogen | Cat# A-21311 |
| **Chemicals, Peptides, and Recombinant Proteins** | | |
| Tissue-Tek O.C.T. Compound | Electron Microscopy Sciences | Cat# 62550-01 |
| BD Cytofix fixation buffer | BD Biosciences | Cat# 554655 |
| PBS (pH 7.4) | Caisson Labs | Cat# PBL06-6X500ML |
| Triton-X-100 | Sigma-Aldrich | Cat# T-9284 |
| Bovine Serum Albumin | Sigma-Aldrich | Cat# A9576-50ML |
| Normal Mouse Serum | Jackson Laboratories | Cat# 015-000-120 |
| Tris buffer (1 M dilute to 0.1M) | Fisher Scientific | BP1756500 |
| Mix-n-Stain CF Dye Antibody Labeling Kits | Biotium | Cat# 92433-92339 |
| N-methylacetamide | Sigma-Aldrich | Cat# M26305 |
| Histodenz | Sigma-Aldrich | Cat# D2158 |
| 1-Thioglycerol | Sigma-Aldrich | Cat# M1753 |
| High-vacuum grease | VWR | Cat# DOWC1597418 |
| Agarose | Fisher Scientific | Cat# 16500500 |
| Immersion Oil, type F | Fisher Scientific | Cat# NC0586121 |
| EndoFit Ovalbumin 100mg | Invivogen | Cat# vac-nova-100 |
| Alhydrogel adjuvant 2% , 250mL | Invivogen | Cat# vac-alu-250 |
| Sucrose, ultrapure DNAse- and RNAse-free | VWR | Cat# 97061-432 |
| **Software** | | |
| CytoMAP | This Manuscript | https://gitlab.com/gernerlab/cytomap |
| Imaris extensions | This Manuscript | https://gitlab.com/gernerlab/imarisxt_histocytometry |
| Imaris | Bitplane | https://imaris.oxinst.com/ |
| LASX | Leica Microsystems | https://www.leica-microsystems.com/products/microscope-software/p/leica-las-x-ls/ |
| FlowJo | FlowJo, LLC | https://www.flowjo.com/ |
| Prism | GraphPad Software | https://www.graphpad.com/scientific-software/prism/ |
| MATLAB | The MathWorks, Inc. | https://www.mathworks.com/products/matlab.html?s_tid=hp_products_matlab |
| **Other** | | |
| UltraComp eBeads Compensation Beads | Fisher Scientific | Cat # 01-2222-42 |
| PAP pen | Vector Laboratories | Cat# H-4000 |

### Table S2 Channel Arithmetic

| Channel name/Sample | Description | Equation |
|---|---|---|
| **Fi. 2** | | |
| All Cell Composite | CD3 + 0.5*CD45.2 + MHC-II | (ch1 .* 0.5) + (ch3.* 0.5) + ch5 |
| **Fig. 5** | | |
| clean-CD31 | corrected spillover from CD11c | (ch7-ch6).*(ch7>ch6) |
| clean-SIRPa | corrected spillover from CD31 | (ch2-ch7).*(ch2>ch7) |
| clean-CD169 | exclude CD207+ subtract CD31 | (ch4-ch13).*(ch4>ch13).*(ch3<70) |
| clean-CD11c | corrected spillover from CD31 | (ch6-ch13).*(ch6>ch13) |
| clean-MHC-II | exclude B220 | (ch8-ch11).*(ch8>ch11) |

| | | |
|---|---|---|
| Composite Myeloid | clean-SIRPa + CD207 + clean-CD169 + clean-CD11c + clean-MHC-II + Clec9a | (ch14.*0.75) + (ch3.*0.75) + (ch15.*0.5) + (ch16.*0.9) + (ch17.*0.75) + (ch9 .* 0.75 .*(ch9>50)) |
| **Fig. 6 Experiment #1** | | |
| Clean CD31 | CD31-0.1*Lyvel | (ch1-0.1.*ch8).*(ch1>0.1.*ch8) |
| Composite Myeloid | CD11c + Clec9a + CD207 + CD169 + SIRPa + CD301b + MHC2 | ch2+ch4+ch5+ch6+ch7+ch9+(ch10.*4) |
| **Fig. 6 Experiment #2** | | |
| Composite Myeloid | CD169 + 2*SIRPa + 2.5*MHC-II(B220<55) + Clec9a+CD207 + 1.5*CD11c - B220 - CD3 + min("") | ((1.5.*ch1.*(ch1>5) + 3.8.*ch4.*(ch4>5) + 2.5.*ch6.*(ch6>5).*(ch5<40) +2.*ch7.*(ch7>5) + ch9.*(ch9>5) + 2.*ch10.*(ch10>5)-0.5.*ch3-0.5.*ch5) +min(min((1.5.*ch1.*(ch1>5) + 3.8.*ch4.*(ch4>5) + 2.5.*ch6.*(ch6>5).*(ch5<40) +2.*ch7.*(ch7>5) + ch9.*(ch9>5) + 2.*ch10.*(ch10>5)-0.5.*ch3-0.5.*ch5)))./2 |
| **Fig. 3** | | |
| Composite Lymphocyte | CD3+2*B220 | ch10+(2.*ch8) |
| Composite Myeloid | CD11c+MHC-II+1.3*CD64 | ch2+ch4+(1.3.*ch9) |
| **Fig. 4** | | |
| Composite Surface Markers | [(CD11c>14)/175 + (CD11b>10.5)/160 + (B220>15)/130 + (CD3>30)/170 + (MHC-II>20)/165]*150 | (ch2.*(ch2>14)./175 + ch3.*(ch3>10.5)./160 + ch4.*(ch4>15)./130 + ch5.*(ch5>30)./170 + ch11.*(ch11>20)./165).*150 |
| Nuclei | Jojo1 – Composite Surface Markers | ch8 - ch13 |

Table S3. Surface creation parameters

| Surface Spots Name | Source Channel | Smoothing: Surface Detail (μm) | Background Subtraction: Diameter of Largest sphere | Absolute Intensity Threshold | Split Touching Objects: Split seed diameter (μm) | Quality Threshold | Voxel Number Threshold | Sphericity Threshold | Diameter (Spots) |
|---|---|---|---|---|---|---|---|---|---|
| **Fig. 2** | | | | | | | | | |
| Surface_1 | All Cell Composite | 20 | | 8 | | | | | |
| Surface_2 | Inverted Ch | 0.655 | | 107 | 3.28 | 8.28 | 10-2781 | 0.583 | |
| **Fig. 5** | | | | | | | | | |
| Myeloid Surfaces | Composite Myeloid | 0.758 | 25 | 7-22 (Sample Dependent) | 7.5-10 (Sample Dependent) | 4.5 | 300 | | |
| All Spots | B220 | | | | | 13.1 | | | 3 |
| **Fig. 6 Experiment #1** | | | | | | | | | |
| Myeloid Surfaces | Composite Myeloid | 1.5 | 12 | 20 | 12 | 10 | 50 | | |
| CD31 Surfaces | Clean CD31 | 2.5 | 15 | 2.25 | 4 | 3.5 | 150 | | |
| Lyve1 Spots | Lyve1 | | true | | | 2 | | | 5 |
| B220 Spots | B220 | | true LN1 (false LN2) | | | 1.5 LN1 (7.5 LN2) | | | 5 |
| **Fig. 6 Experiment #2** | | | | | | | | | |
| Myeloid Surfaces | Composite Myeloid | 0.8 | 11 | 40 | 10 | 6 | 50 | | |
| CD31 Surfaces | CD31 | 2 | 5 | 2 | 4 | 2.5 | 50 | | |
| Lyve1 Spots | Lyve1 | | false | | | 20 | | | 5 |
| B220 Spots | B220 | | false | | | 40 | | | 5 |
| **Fig. 3** | | | | | | | | | |
| Myeloid Surfaces | Composite Myeloid | 0.6 | 11 | 20 | 11 | 5 | 50-7103 | | |
| Lymphocyte Surfaces | Composite Lymphocyte | 0.6 | 10 | 9 | 6 | 2.5 | 50 | | |
| **Fig. 4** | | | | | | | | | |
| All Cells | Nuclei | 0.63 | 10 | 18.5 | 3.16 | 6.08 | 10 | | |

Table S4. Pseudo-space input parameters

| |
|---|
| Fig. 2f<br>Data Preparation: Neighborhood Composition |

| Cell Type | Weight | Smoothing |
|---|---|---|
| B cells | -1 | 500 |
| T Cells | 1 | 500 |
| DCs | 0.2 | 500 |
| Fig. 3e<br>Data Preparation: Number of cells per Neighborhood | | |
| Type | Weight | Smoothing |
| B cells | 1 | 5000 |
| Teff | 0 | 5000 |
| Treg | 5 | 5000 |
| TAM | -2 | 5000 |
| DCs | 0 | 5000 |
| aMACs. | 1 | 5000 |
| SIRPα$^{DIM}$MHC-II+ | -1 | 5000 |
| Fig. 4e<br>Data Preparation: Number of cells per Neighborhood | | |
| B Cells | 1 | 1000 |
| Alv. Macs | -50 | 1000 |
| CD11b+ | 0 | 500 |
| DCs | 0 | 1000 |
| Mtb+ Cells | 100 | 100 |
| T Cells CD4- | 1 | 1000 |
| T Cells CD4+ | 1 | 1000 |

### Table S5. Antibody staining panels and Microscope Settings

| Ch | Antibody | Flurophore |
|---|---|---|
| **Fig. 2** | | |
| 1 | CD3 | 421 |
| 2 | TCF1 | PacBlue |
| 3 | CD45.2 (OT-II) | 510 |
| 4 | CXCR3 | PE |
| 5 | MHC-II | 594 |
| 6 | Ki67 | 700 |
| 7 | B220 | CF750 |
| 8 | anti-GFP | AF488 |
| 9 | CD11c | 647 |
| **Fig. 5** | | |
| 1 | CD64 | PE |
| 2 | SIRPα | 594 |
| 3 | CD207 | 488 |
| 4 | CD169 | 514 |
| 5 | Lyve1 | 490LS |
| 6 | CD11c | 421 |
| 7 | CD31 | 480 |
| 8 | MHC-II | 395xl |
| 9 | Clec9a | 633 |
| 10 | CD301b | 660 |
| 11 | B220 | 700 |
| 12 | CD3 | APC-f750 |
| **Fig. 6 Experiment #1** | | |
| 1 | CD31 | BV421 |
| 2 | CD11c | BV480 |
| 3 | B220 | Dy405ls |
| 4 | Clec9a | CF633 |
| 5 | CD207 | AF488 |
| 6 | CD169 | CF514 |
| 7 | SIRPa | CF594 |
| 8 | Lyve1 | eFluor570 |
| 9 | CD301b | CF660 |
| 10 | MHC-II | AF700 |
| **Fig. 6 Experiment #2** | | |
| 1 | CD169 | CF514 |
| 2 | CD3 | AF700 |
| 3 | Lyve1-biotin | SA-CF750 |
| 4 | SIRPa | BV421 |
| 5 | B220 | BV510 |
| 6 | MHC-II | DY395XL |

| | | |
|---|---|---|
| 7 | Clec9a | CF633 |
| 8 | CD31 | AF488 |
| 9 | CD207 | AF546 |
| 10 | CD11c | eFluor615 |
| **Fig. 3** | | |
| 1 | SIRPa | BV421 |
| 2 | CD11c | BV480 |
| 3 | CD8 | BV510 |
| 4 | MHC-II | Dy396XL |
| 5 | Clec9a | αsheep-CF633 |
| 6 | PCREB | AF488 |
| 7 | IRF4 | CF514 |
| 8 | B220 | SA-490ls |
| 9 | CD64 | CF660c |
| 10 | CD3 | AF700 |
| 11 | PD1 | αgoat-CF750 |
| 12 | Foxp3 | eF570 |
| 13 | PD-L1 | CF594 |
| **Fig. 4** | | |
| 1 | iNOS | AF405 |
| 2 | CD11c | BV480 |
| 3 | CD11b | |
| 4 | B220 | 405ls |
| 5 | CD3 | CF633 |
| 6 | pS6 | AF488 |
| 7 | IFNg | PE |
| 8 | Jojo1 | |
| 9 | IRF4 | CF594 |
| 10 | CD4 | CF660 |
| 11 | MHC-II | AF700 |
| 12 | Mtb | CF750 |
